# MALE-MALE SOCIAL BONDING, COALITIONARY SUPPORT AND REPRODUCTIVE SUCCESS IN WILD GUINEA BABOONS

**DOI:** 10.1101/2022.04.19.488751

**Authors:** Federica Dal Pesco, Franziska Trede, Dietmar Zinner, Julia Fischer

## Abstract

Male-male bonds may confer substantial fitness benefits. The adaptive value of these relationships is often attributed to coalitionary support, which aids in rank ascension and female defence, ultimately resulting in greater reproductive success. We investigated the link between male-male sociality and both coalitionary support and reproductive success in wild Guinea baboons. This species lives in a tolerant multi-level society with reproductive units comprising a male and 1-6 females at the core. Males are philopatric, form differentiated, stable, and equitable affiliative relationships (‘strong bonds’) with other males, and lack a clear rank hierarchy. Here, we analysed behavioural and paternity data for 30 males and 50 infants collected over four years in the Niokolo-Koba National Park, Senegal. Strongly bonded males supported each other more frequently during conflicts, but strong bonds did not promote reproductive success. Instead, males that spent less time socializing with other males were associated with a higher number of females and sired more offspring. Notably, reproductively active males still maintained bonds with other males, but adjusted their social investment in relation to life-history stage. Long-term data will be needed to test if the adaptive value of male bonding lies in longer male tenure and/or in promoting group cohesion.

## Introduction

According to sexual selection theory [1,2], males with higher quality should have greater reproductive success. In numerous species, males with the best fighting ability, i.e., the greatest strength or the best weapons, have advantages in male-male competition, gain higher dominance ranks and better access to fertile females, and sire the highest number of offspring [3]. A classic case are Northern elephant seals (*Mirounga angustirostris*), where the heaviest males reap the vast majority of matings [4]. Such intrasexual competition is typically more distinct in males, whereas mate choice is more prevalent in females [5,6]. Females may prefer males that have more exaggerated ornaments [1,7] or that spend more time and energy in elaborate courtship displays [8]. In group living animals, male reproductive success may not only depend on strength or ‘beauty’, but also on ‘social capital’, that is, the ability to cooperate and forge bonds with other males.

As observed in a wide range of taxa, including non-human primates, lions, horses, dolphins, and some species of bird and fish, cooperation between males can aid in female defense resulting in longer tenure and/or increased number of females and offspring [9-15]. A prime example are male lions (*Panthera leo*) where larger coalitions are more successful in taking over female prides resulting in longer tenure and greater number of surviving offspring [9]. Similar mechanisms occur in some multi-level primate societies, where ‘leader males’ with associated ‘follower males’ have longer tenure, higher numbers of females, and more offspring [12,13].

Enhanced reproductive success has also been linked to ‘strong bonds’ between males, defined as affiliative relationships that are differentiated, equitable, and stable over time [16]. A number of studies have shown that investments in strong bonds are linked to increased coalitionary support, which in turn results in rank ascension and, ultimately, enhanced reproductive success [17-19]. In Chimpanzees (*Pan troglodytes*) males siring success is also associated to the establishment of a large network of strong ties with other males [18]. In addition, male-male bonds may also affect female choice, as male coalitions may reduce harassment from other males and decrease infanticide risk [10,12,20] or offer better protection against predators [21].

We investigated the reproductive benefits of strong bonds between males in wild Guinea baboons (*Papio papio*). Guinea baboons live in a nested multi-level society, with ‘units’ composed of a ‘primary’ male, one to six associated females, and immatures at the core of the society [22]. Several units, together with ‘bachelor’ males, make up a ‘party’ and two to three parties regularly aggregate into ‘gangs’ with overlapping home ranges [23]. Most primary males (76.5%) have one or more associated bachelors and bachelor males are often (66.7%) associated to multiple units [24]. Primary males maintain largely exclusive affiliative and sexual relationships with the females in their unit, while bachelors exchange a smaller proportion of social interactions with females and are usually not reproductively active [25]. ‘Solitary’ males, as observed in hamadryas baboons [26], occur only rarely [24]. Males are predominately philopatric, display a high degree of spatial tolerance, form strong bonds, and support each other in coalitions [23,24]. Strongly bonded males are on average more closely related than less strongly bonded males indicating that kin selection plays a role in male-male bonding [24]. Nevertheless, relatedness does not seem to fully explain male-male relationship patterns in our study population [22]. Compared to other baboon species, males show low rates of aggression and no clear dominance hierarchy [24], while females have high levels of spatial freedom and play an active role in the formation and maintenance of inter-sexual relationships [25].

We predicted that strong bonds between males - enhanced by coalitionary support - would result in higher male reproductive success via the attraction of more females, resulting in a higher number of offspring. To test this prediction, we determined bond strength following Dal Pesco et al. [24] and assessed the link between bond strength and coalitionary support. We predicted that dyads with stronger bonds would be more likely to cooperate during conflicts. Our core analysis examined whether male bond strength and the number of strong bonds a male has were linked to enhanced reproductive success in the form of increased numbers of associated females and sired offspring.

## Materials and Methods

### Field site, study subjects, and data collection

Throughout the course of 45 months (April 2014 - December 2017), we collected data on wild Guinea baboons - one of six baboon species [27] - at the Centre de Recherche de Primatologie (CRP) Simenti field station in the Niokolo-Koba National Park in Senegal (described in [22]). During the study period, the Simenti Guinea baboon community comprised ∼ 400 individuals including five habituated parties in two gangs. The two parties with the highest number of males were selected as our study groups (party 9 from the Mare gang and party 6 from the Simenti gang). We used the party as our group unit and restricted all analyses to within-party interactions [24,28]. Party size and composition varied during the study period due to maturation, dispersal/migration, and mortality with an average of 43 individuals in party 6 (range: 35-48, average adult sex ratio (male:female) of 1.03) and 46 individuals in party 9 (range: 38-51, average adult sex ratio (male:female) of 0.48).

We performed behavioral observations of all adult and all small and large subadult males belonging to the two study parties (n=30; party 6, n=16; party 9, n=14). Males were included as focal subjects when they were first classified as small subadult (∼ 6 years old). At this age, they already establish close affiliations and display strong bonds and coalitionary support [24,29] with adult males. All details about male presence and age category changes, age category assessment, and criteria for subject selection/exclusion can be found in supplementary appendix 1, tables S1 and S2, and figure S1. We conducted behavioral observations for a total of 872 observation days (1956 contact hours for party 6 and 1954 contact hours for party 9). All data were collected using customized electronic forms developed for our long-term data collection in the Pendragon 7.2 software (Pendragon Software Corporation, USA) on Samsung Note 2 handhelds. We recorded census information about demographic changes (including birth, death, dispersal/migration, presence/absence), health status, and female reproductive state [25] on every observation day. In all analyses, we controlled for the time a male was present in the study party, due to entering the sub-adult age category or death/disappearance.

We conducted 20-mins focal follows [30] balanced between subjects and time of day, for an average of five monthly protocols per individual and a total focal time of 1547 h (total number of focal protocols = 4787). Protocols included recordings of continuous focal animal activity (i.e., moving, feeding, resting, and socializing) and all occurrences of social behaviors such as approach within 1 m, retreat, grooming, contact-sit, and greeting. All grooming and contact-sit durations were recorded to the nearest second. Instances of aggression, coalitionary support, copulation and grooming were additionally recorded ad libitum. Coalitionary support was scored every time two or more individuals simultaneously directed aggression toward a common target that could be a single male or another male-male coalition. Only coalitions involving two male allies against a common male target were included in our analysis. Due to the very low rate of aggression, all occurrences of coalitionary support between males, including both focal and ad libitum events, were included in our analysis.

### Male-male social bonds and unit composition

We used the dyadic composite sociality index (hereafter DSI [31]) to quantify dyadic affiliative relationship strength. This index ranges from 0 to infinity with a mean of 1, and measures the deviation of affiliative behavior of a given dyad compared to all other dyads in the same group. The DSI is calculated using the following formula: 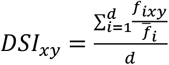 where *f*_*ixy*_ is the behavioral rate for dyad *xy* and behavior *i*, 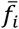 is the average behavioral rate for behavior *i* calculated across all dyads in the party, and *d* is the number of behaviors included in the index calculation [31]. We computed yearly DSI values (January to December) for each male-male dyad within the party using the following positively correlated affiliative behaviors: grooming frequency and duration, contact-sit frequency and duration, and frequency of within 1 m approaches [24]. To avoid redundancies with other behaviors, only approaches that were not followed by social behavior (positive or negative) within 10 s were considered in the DSI calculation. Individual bond strength was calculated as the sum of a male’s top three OSI values [32]. The number of strong bonds per male was based on the number of higher than average DSI values [33].

Data on female-male interactions (i.e., frequency of copulations, grooming bouts, contact-sit bouts, greetings, and aggression events and duration of grooming and contact-sit bouts), unit composition, and female unit transfers were recorded on every observation day. Following established methodologies [24,28] based on previous findings showing that females exchange significantly higher rates of interactions with their primary male [25], we used female-male interaction occurrence to verify daily unit composition within each study party.

### Genotyping and paternity analysis

To establish paternity, we collected fecal samples of all subadult and adult males (n=30) and subadult and adult females (n=33) in party 6 and party 9. Fifty infants were born during the study period in these two parties. We were able to collect fecal samples from 36 infants for paternity analysis, while the remaining 14 infants deceased before sampling could occur. To check for extra-party paternities, we additionally sampled all subadult and adult males (n=17) belonging to the other three habituated parties of our study population as well as two adult males that were associated with party 6 for only 36 days (see supplementary appendix 1).

We evaluated individual allelic variation based on 24 polymorphic autosomal microsatellite markers. This microsatellite panel [34] is an optimized version of the panel that was successfully used in several studies of Guinea baboons [e.g. 35] and our own study population [23]. Genetic sample collection, storage, DNA-extraction, and genotyping methodologies are described in detail in Dal Pesco et al. [34]. Detailed information about number and type of samples available per individual can be found in supplementary appendix 2 and 3.

Following the methodologies in Dal Pesco et al. [34], we calculated descriptive statistics for all 24 markers (including F_IS_, expected and observed heterozygosity) and tested for Hardy-Weinberg equilibrium (HWE) and presence of null alleles (see table S3). All loci were polymorphic with allele numbers averaging 4.0 (SD=1.4, range=2.0 to 7.0). As locus D1s548 showed signs of null alleles, it was excluded from the paternity analysis, which was therefore performed using a total of 23 loci.

We estimated paternity using the software Cervus (version 3.0.7) [36] and following the methodologies explained in detail in Dal Pesco et al. [34]. We recorded the identity of the mother during field observations, and additionally checked all mother/offspring pairs with a maternity likelihood analysis (criteria for acceptance: identification as candidates with 0 mismatches). All mothers were confirmed with 0 mismatches. We then used a trio likelihood approach where the identity of the mother was known to determine the most likely father (see table S4). A male was considered to have sired an offspring when he was assigned as the most likely father, had a maximum of 1 mismatched allele, and the confidence level for the assignment was more than 95% (‘strict’ criterion).

### Statistical analyses and modeling

All statistical analyses and figure preparation were performed in the R environment (version 4.0.5) [37] using the RStudio interface (version 1.4.1106-5) [38]. We ran Generalized Linear Mixed Models (GLMM) [39] using the R packages ‘lme4’ (version 1.1-26) [40] for all Poisson models and ‘glmmTMB’ (version 1.0.2.9000) [41] for the beta model used in our post-hoc analysis.

To reduce type I error rates, we used the maximal random-effect structure comprising all theoretically identifiable random slope components [42] excluding the correlations between random intercepts and slopes when “unidentifiable” (i.e., absolute correlation parameter ∼1) [43]. To ease model conversion and estimate comparison, all covariates were z-transformed to a mean of zero and a standard deviation of one prior to fitting each model [44]. Detailed information about sample size, model complexity, checks for the need of zero inflation, random slopes, data standardization/transformation (including means and standard deviations of original values), model stability, and the use of non-default optimizers can be found in the supplementary appendix 4 and the tables S5-S10.

Before inference, all models were validated using diagnostic checks. We assessed the absence of collinearity among predictors calculating the Variance Inflation Factors (VIF) [45] using the ‘vif’ function of the package car (version 3.0-10) [46] on reduced general linear models with all random effect structures and optimizers excluded. With an overall maximum VIF value of 1.95 we ruled out collinearity for all our models. We evaluated the assumption of normality for each random effect component by visually inspecting histograms of each random intercept and slope. No obvious deviation from these assumptions was recorded. For all models (see details in each sub-section) we calculated the dispersion parameter to check for potential type I errors due to overdispersion.

In models with multiple predictors of interest, we first determined the significance of the full model (also including all predictors of interest) against a null model comprising only the control predictors and the random-effect structure using a likelihood ratio test [47]. This allowed us to test the overall effect of our predictors of interest avoiding ‘cryptic multiple testing’ [48]. P-values for individual predictors were obtained using the likelihood ratio test of the ‘drop1’ R function with argument ‘test’ set to ‘chisq’ [42]. The function ‘bootMer’ of the package ‘lme4’ was used to perform a parametric bootstrap (1000 bootstraps) and obtain 95% model estimate confidence intervals. Effect sizes were calculated using the ‘r.squaredGLMM’ function of the ‘MuMIn’ R package (method Trigamma; version 1.43.17) [49] and the ‘r2’ function of the ‘performance’ R package (version 0.7.1) [50] for all Poisson models and the beta model used in our post-hoc analysis, respectively.

### Male-male sociality and coalitionary support

To investigate whether males with stronger bonds were more likely to support each other in coalitions we ran a GLMM [39] with Poisson error structure and log link function [51] where dyadic coalitionary support frequency per year (including both focal and ad libitum events) was the count response and yearly DSI (log- and then z-transformed) was the predictor of interest. To control for observation effort, we included the log-transformed contact time in hours as an offset term [51]. Note that contact time was calculated using the total time spent working with each study party during each daily working session and taking into account demographic changes for each male-male dyad. We included year and party membership as fixed control factors, and male identities (subject identity and coalition partner identity) and dyad identity (composed by subject identity followed by coalition partner identity) as random intercepts. The following random slope components were also included: year (manually dummy coded and then centered) and DSI (z-transformed) within both male identities. The model was not overdispersed (dispersion parameter= 0.289).

### Male-male sociality and reproductive success

To examine if greater levels of male-male sociality were associated with enhanced male reproductive success we analyzed two different measures of male reproductive success: number of associated females and number of sired offspring. To account for unit size variation due to female transfers and demographic changes, we used daily unit size data to calculate the number of associated females within each year as a yearly mode per male (i.e., the most frequent unit size value). The number of sired offspring was calculated as the sum of sired offspring per male within each year (n=49; one offspring was fathered by a male of another party; see table S4). As within each unit paternity probability for the primary male is very high (for this dataset 91.7% of offspring were sired by the mother’s primary male at time of conception), for the 14 offspring for whom we had no genetic data, we selected the mother’s primary male at the time of conception as the most likely father. Our measures of male-male sociality were male bond strength, calculated as the yearly sum of a male’s top three OSI values, and number of strong bonds, calculated as the yearly number of higher-than-average DSI values per male.

We ran two GLMMs [39] with Poisson error structure and log link function [51], where the yearly mode of number of associated females and yearly number of sired offspring were the count responses and male bond strength and number of strong bonds were the predictors of interest. In both models, we included year and party membership as fixed control factors, and male identity as random intercept. The following random slope component was also included in both models: male bond strength (z-transformed) within male identity. Both models were not overdispersed (dispersion parameters=0.542 and 1.159).

### Post-hoc analysis: time males spent affiliating with other males by number of associated females

In light of the results of our analysis, we performed a post-hoc investigation to look at the effect of the number of associated females on the proportion of time males spent affiliating (i.e., grooming and contact-sit) with other males. This allowed us to specifically look at male time budgets and to analyze interaction occurrence, which can in some cases represent social relationships more accurately and precisely compared to composite sociality indices [52]. We ran a GLMM [39] with a beta error structure and logit link function [51,53] with the proportion of time males spent affiliating with other males as the response and number of associated females as the predictor of interest. To avoid response values being exactly zero or one, we transformed the response prior to fitting the model using the following formula x^1^=(x*(length(x)-1)+0.5)/length(x) [54]. We included year and party membership as fixed control factors, and male identity as random intercept. The model presented signs of moderate overdispersion (dispersion parameters=1.283), which could not be resolved by specifically modelling dispersion with the argument ‘dispformula’. To adjust for overdispersion and the increase type I error rate, we corrected the estimate standard errors by the overdispersion level according to Gelman and Hill (SEadjusted= SE x ✓dispersion parameter) [55]. Furthermore, z-and p-values were determined again based on the adjusted standard error with z=estimate/ SEadjusted and p=2*pnorm(q=-abs(z)).

## Results

### Male-male sociality and coalitionary support

Males maintained differentiated male-male relationships, with DSI values ranging from 0.00 to 21.03 (SD=2.29; median=0.06). About a fifth (20.7%) of the dyads had a DSI above the party average. The average bond strength per male was 9.35 (SD=6.51; range=0.27 to 33.95) and the average number of strong bonds per male was 2.18 (SD=1.52; range=0 to 6). The average DSI across all strongly bonded male dyads was 4.37, indicating that these dyads affiliated four times as often/long as compared to the average of the party.

A total of 290 two-against-one coalitions were recorded between males during the study duration (both from focal and ad libitum data) with 26.9% of dyads (n=53 of 197) engaging in at least one coalition. Overall, dyads supported each other on average 1.47 times (SD=4.78; range=0 to 36) across the study period with an average rate per hour of 0.001 (SD=0.003; range=0.000 to 0.021) coalitions. Dyads with higher DSI values were more likely to support each other in coalitions (estimate ± SE=0.781±0.108, CI[0.500,0.994], p<0.001, figure 1, also see table S5).

**Figure 1.**
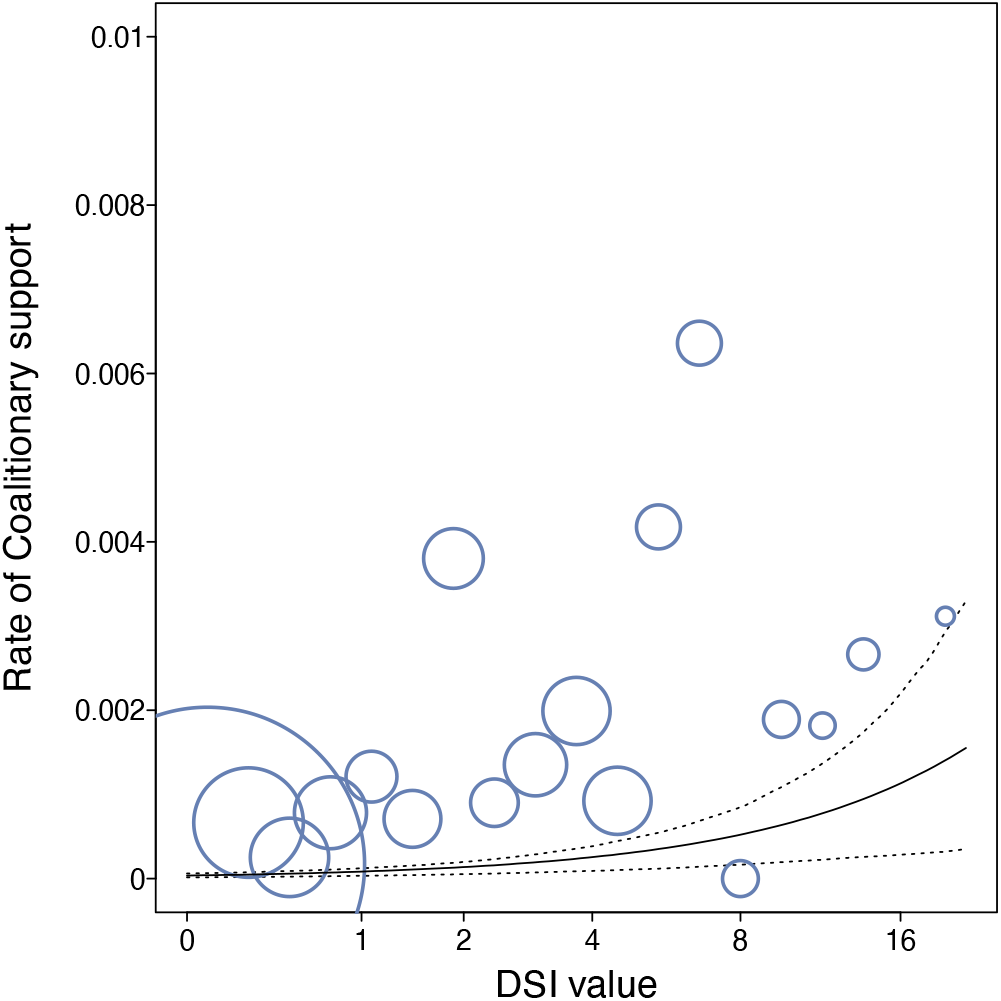
Relationship between male-male dyadic bond strength (DSI value) and dyadic rate of coalitionary support. Dyads with stronger bonds were more likely to support each other in coalitions (GLMM: n=958, p<0.001). DSI values are represented in log-scale and binned in 19 bins. The area of the circles depicts the frequency with which a given number of coalitions per contact hour occurred in a given bin (mean=3.29, range=1 to 232). The solid line depicts the fitted model and the dashed lines depict the bootstrapped 95% confidence intervals with all other predictors being at their average (party and year manually dummy coded and centered).

### Male-male sociality and reproductive success

Twenty-one of the 30 study males had at least one associated female during part or the entire study period, while the remaining males were not associated with a female during the study period (see figure 2). Of the nine males that never had primary status, seven were subadult males and two were old adult males for most of the study time during which they were present in the study party, corroborating the observation that bachelor males are often subadult or late-prime/old males [24]. Of the 21 males that had primary status at least once, twelve were adult males for their entire presence time, eight transitioned from subadult to adult during the study period, and one was a large subadult male during his presence time (see figure S1 for male age category changes).

**Figure 2.**
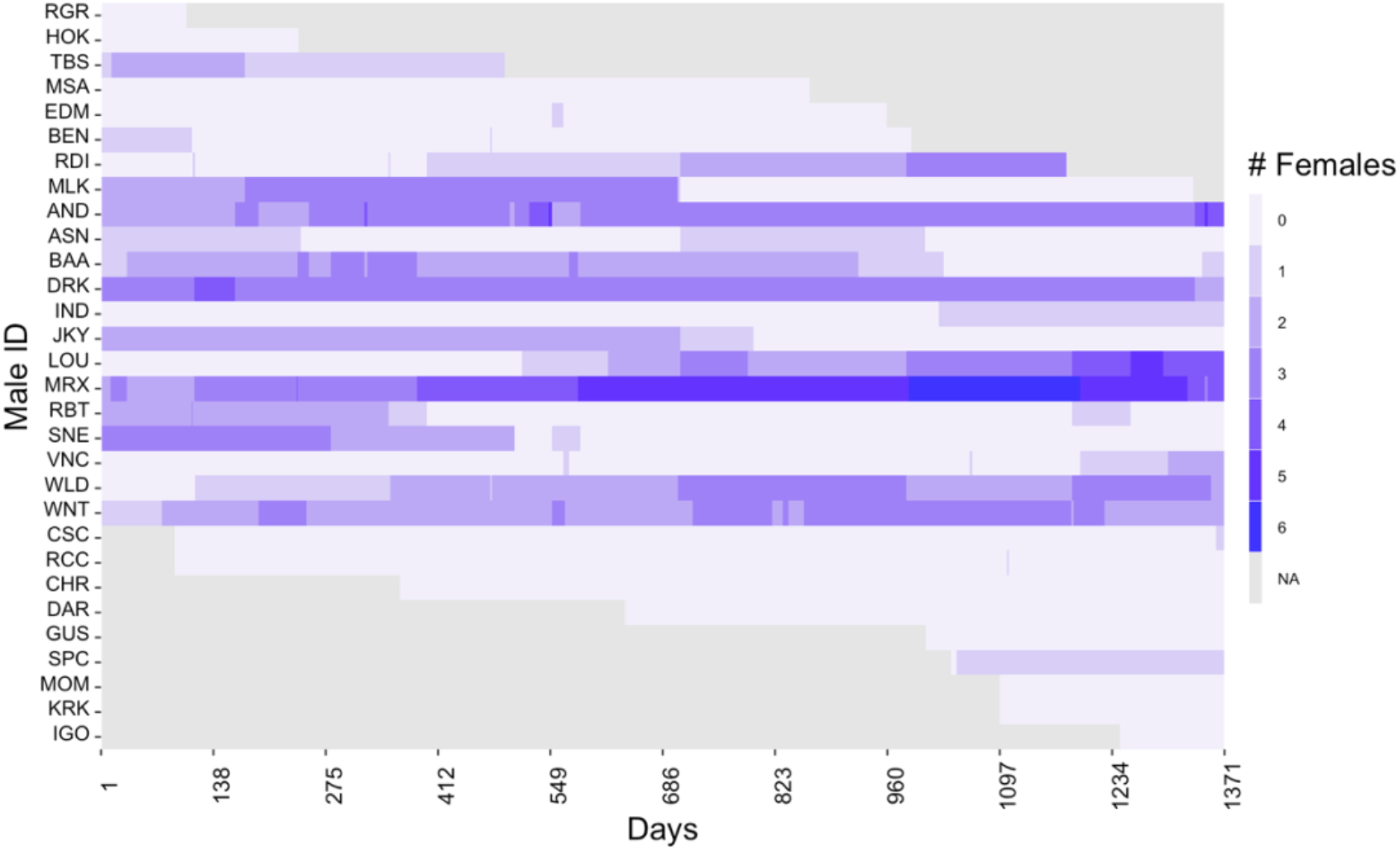
Visualization of the variation in male status and unit size (i.e., number of associated females) over the course of the study period (April 2014 - December 2017) for the 30 study subjects. NA (not assessed - in grey) indicates days when males were not present due to demographical changes (i.e., not associated to the study parties, not in the selected age category, or deceased, also see supplementary appendix 1 and figure S1).

The average mode of the number of associated females per male per year was 1.09 (SD=1.40; range=0 to 6). This average was 1.29 (SD=1.43; range=0 to 6) if we excluded males that never had primary status throughout the study period. The full model including the two predictors of interest (male bond strength and number of strong bonds) accounted for significantly more variance compared to the null model (full null model comparison: *χ*^*2*^=22.237, p<0.001). While there was no obvious evidence that the number of strong bonds had an effect on the number of associated females (estimate±SE=-0.288±0.220, CI[-0.779,0.176], p=0.181), we found strong evidence that males with higher bond strength were associated with fewer females (estimate± SE=-0.749±0.266, CI[-1.381,-0.254], p=0.003; figure 3a and 4a, also see table S6). The negative effect of male bond strength held true when we analyzed a subset of data only including adult and large subadult males within each year (estimate± SE=-0.739±0.259, CI[-1.368,-0.281], p=0.002; see table S9).

**Figure 3.**
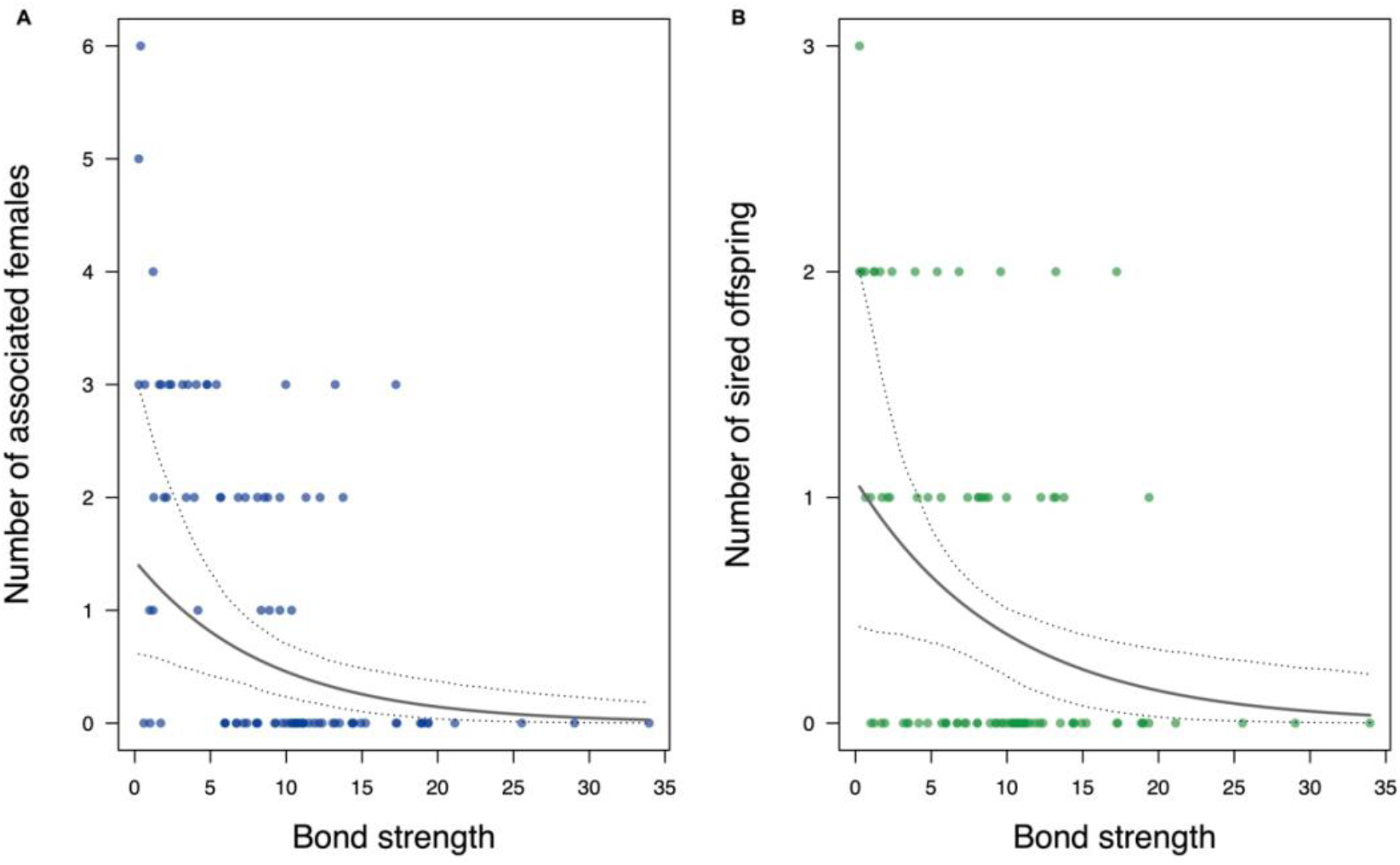
Relationship between male bond strength (calculated as the sum of a male’s top three D51 values) and A) number of associated females (mode per male per year) and B) number of sired offspring (count per male per year). Males with stronger bonds were found to have fewer associated females (GLMM: n=91, p=0.003) and to sire fewer offspring (GLMM: n=91, p=0.017). Points represent each subject in a given year (2014-2017). The solid line depicts the fitted model and the dashed lines the bootstrapped 95% confidence intervals with all other predictors being at their average (party and year manually dummy coded and centered and number of strong bonds z-transformed to a mean of 0 and standard deviation of 1).

**Fig 4.**
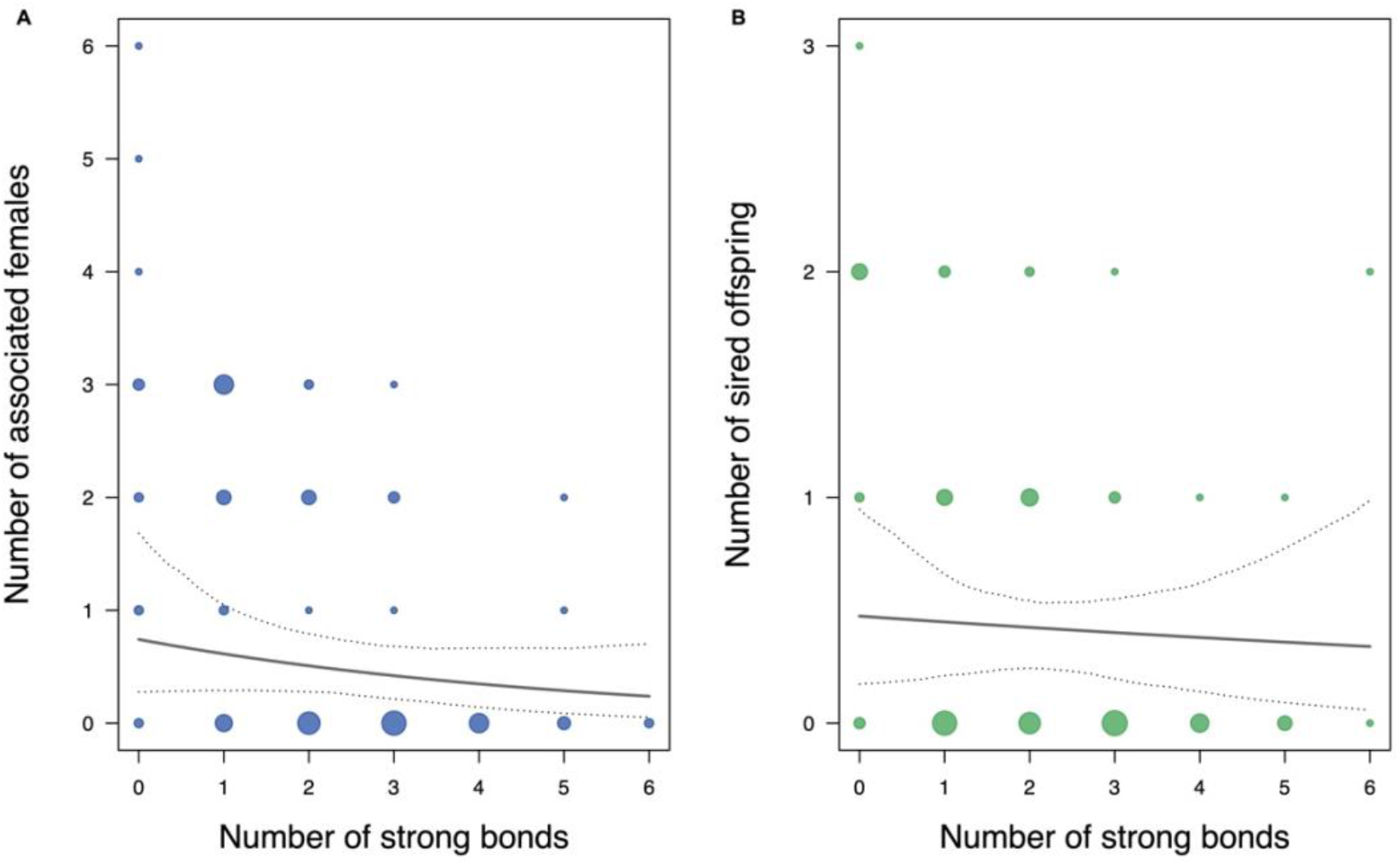
Relationship between number of strong bonds (calculated as the number of higher-than-average DSI values per male) and A) number of associated females (mode per male per year) and B) number of sired offspring (count per male per year). There was no evidence for a relationship between number of strong bonds and number of associated females (GLMM:, n=91, p=0.181) or number of sired offspring (GLMM: n=91, p=0.727). Points represent each subject in a given year (2014-2017). The solid line depicts the fitted model and the dashed lines the bootstrapped 95% confidence intervals with all other predictors being at their average (party and year manually dummy coded and centered and number of strong bonds z-transformed to a mean of 0 and standard deviation of 1).

Overall males in the study parties sired 49 offspring (one offspring was fathered by a male of another party) with an average number of 1.63 (SD=2.13; range=0 to 8) offspring sired per male across the study period. The average number of offspring sired across the study period was 2.33 (SD=2.20; range=0 to 8) if we only considered males that had primary status at some point during the study period. The average number of offspring sired per male per year was 0.54 (SD=0.78; range=0 to 3). This average was 0.64 (SD=0.81; range=0 to 3) if we excluded males that never had primary status throughout the study period. The full model with the two predictors of interest (male bond strength and number of strong bonds) accounted for significantly more variance compared to the null model (full-null model comparison: *χ*^*<sup>2</sup>*^ =11.260, p=0.004). While there was no obvious evidence that number of strong bonds had an effect on the number of sired offspring (estimate±SE=-0.085±0.245, CI[-0.629,0.367], p=0.727), we found moderate evidence that males with higher bond strength sired fewer offspring (estimate± SE=-0.655±0.288, CI[-1.358,-0.144], p=0.017; see figure 3b and 4b; also see table S7). The negative effect of male bond strength held true when we analyzed a subset of data only including adult and large subadult males within each year (estimate± SE=-0.664±0.289, CI[-1.372,-0.151], p=0.013; see table S10).

### Post-hoc analysis: effect of number of associated females on time spent affiliating with other males

Contrary to our predictions, male bond strength was linked to lower numbers of associated females. We therefore performed a post-hoc analysis focused on male time budget to explore the relationship between time spent affiliating with other males and the number of associated females. We found strong evidence that males with higher numbers of associated females spent a lower proportion of time affiliating with other males (estimate±SE=-0.371±0.107, CI[-0.550,-0.209], p<0.001; figure 5; see table S8).

**Figure 5.**
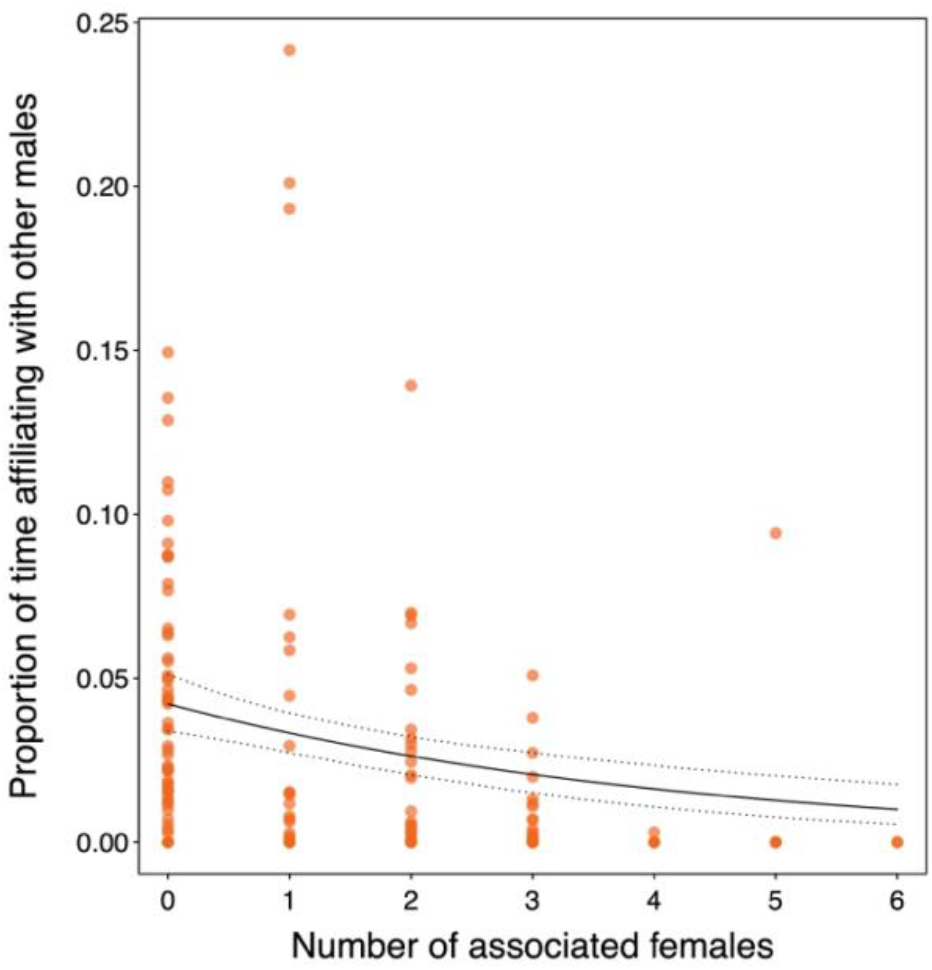
Effect of the number of associated females per male on the proportion of time males spent affiliating with other males. Males with higher numbers of associated females spent lower proportions of time affiliating with other males (GLMM: n=147, p<0.001). Points represent each dyad in a given year (2014-2017). The solid line depicts the fitted model and the dashed lines depict the bootstrapped 95% confidence intervals with all other predictors being at their average (party and year manually dummy coded and centered).

## Discussion

Contrary to our predictions, we found no evidence that male-male sociality was linked to higher reproductive success. Instead, we observed a strong negative relationship between bond strength and male reproductive success (i.e., number of associated females and paternities). Guinea baboon males that were associated with a higher number of females spent less time affiliating with other males. While a number of now classic studies reported a positive relationship between sociality and reproductive success in both males and females across several mammalian [17,18,32,56] and bird species [14], our results indicate that male-male sociality need not directly translate into increased short-term reproductive success. Instead, males that invest time and energy in relationships with females, at the expense of relationships with males, have the highest reproductive success.

Interestingly, reproductive success was not obviously negatively related to the number of strong bonds a male had, indicating that males do maintain differentiated relationships with other males, but mainly adjust their time budgets in relation to the number of females they are able to attract. As intersexual bonding patterns in this species are largely driven by female choice [25] and in light of the high paternity certainty within units (91.7% in this study), stable bonds with females confer direct fitness benefits. It therefore pays for males to invest in bonds with females, irrespective of their reproductive state [25]. Similar patterns were observed in horses (*Equus caballus*), where less successful stallions maintained stable alliances with others, while successful ones exclusively focused on their mares [10].

Guinea baboon males appear to face a trade-off between investments in same-sex and opposite-sex bonds, and the investment in different types of bonds varies with life-history stage: young and old bachelor males invest more in same-sex relationships, but turn their attention to females once they have become primary males - at the expense of time available for their male ‘friends’. Similar effects are seen in male Barbary macaques (*Macaca sylvanus*) and snub-nosed monkeys (*Rhinopithecus bieti*) across seasons, where investment in male-male affiliative relationships drops during the mating season [57,58].

Long-term data will be needed to assess whether male-male bonds are related to an earlier/later acquisition of females, thus increasing tenure length and in this way reproductive success. Additionally, bonds may constitute a “fall-back” option for males once they lose their status as a primary male by providing support and tolerance in old age, and indirectly promote group cohesion. Indeed, Barbary macaque males rely more heavily on cooperative strategies during their post-prime phase [59], while older chimpanzees show greater levels of positive behaviors as well as higher numbers of mutual male-male relationships [60]. For now, we are confident that male-male sociality is negatively linked to reproductive success over the short time, but cannot exclude the possibility that bonds increase life-time reproductive success via earlier or longer male tenure.

How do males manage their relations with other males, when most of their social investments go to females? Under time budget constraints, Guinea baboon males may use male-male ritualized greeting behaviour, characterized by quick, stylized and costly exchanges [28], to assess and maintain their relationships. We propose that the most intense and potentially costly forms of greetings, which occur more often between strongly bonded males [28], can play a central role in male-male bond maintenance once males acquire primary status and invest less time in affiliation. Similarly, in macaques, ritualized interactions between males have been proposed as efficient means in maintaining bonds when their social time with other males is limited [61].

Regarding coalition formation, Guinea baboon males with stronger bonds supported each other more often during agonistic events, corroborating previous analyses in the same [23] and several other species [17,18,57,62]. Compared to macaques, however, rates of coalitionary support in Guinea baboons are low (0.001/hr; Assamese macaques, *Macaca assamensis*: 0.11/hr [17]; Barbary macaques: 0.01-0.21/hr [57]), mirroring the low rate of aggression [24]. Given the lack of a clear dominance hierarchy between males [24] and the presence of frequent instances of coalitions targeting other coalitions [29], it is unlikely that coalitions serve in rank ascension. Why Guinea baboon males engage in possibly risky coalitions and what benefits strong bonds and cooperation may confer requires further investigation.

Ultimately, Guinea baboon females may not gain much from preferring males with strong bonds, as males rarely attempt to takeover females from other males, and infanticide has not been observed in this population [22,25]. Moreover, females do not appear to choose males with strong bonds as means of protection from predators [21]. Instead, females may simply prefer males in good condition. Indeed, mane coloration and length, as well as hind-quarter coloration have been proposed as honest signals of male quality in hamadryas and Guinea baboons [63,64], a hypothesis that remains to be tested. In male geladas, redder chest patches are associated with higher status and larger units [65]. Our current working hypothesis is that male condition and attention to the female are the key determinants of female choice. Although females may have preferences for specific males, female benefits may decrease in larger units due to higher levels of female-female competition over social support and mating opportunities [66]. Female choices are therefore likely also affected by the size and composition of the unit. Considering that males sometimes show parental care (pers. observation) and are generally tolerant toward females, it is also possible that females take into account a male’s willingness to provide care for offspring, as reported in mountain gorillas [67], or a male’s disposition to accept females’ spatial freedom [68]. Long-term data will be needed to test these ideas.

Taken together, we suggest that female choice explains male-female associations, while female-female competition may result in an upper limit on unit size. Consequently, almost all males in their prime achieve some reproductive success and there is little to fight over. Variation in male-male sociality may thus be conceived as an outcome of males adjusting their affiliation patterns according to females’ choices. Our study reinforces the view that male strategies may vary considerably in relation to female leverage in mate choice, and that even among closely related species, such as in the genus *Papio*, entirely different strategies may evolve. Our findings add a piece to the puzzle of understanding the co-evolutionary dynamics of male and female strategies.

## Acknowledgments

We thank the Oirection des Parcs Nationaux and Ministere de l’Environnement et de la Protéction de la Nature de la République du Sénégal for permission to work in the Niokolo Koba National Park. We particularly thank: the former conservateurs of the park Colonel Ousmane Kane and Commandant Mallé Gueye for their cooperation and logistical support; all the staff and field assistants of the CRP Simenti - in particular Mustapha Faye, Armél Louis Nyafouna, Elhadji Dansokho, Lamine Diedhiou, Moustapha Dieng and Touradou Sonko, Matthis Drolet, Lauriane Faraut, Mathieu Cantat, Fransziska Wegdell, Eréna Dupont - for the for their assistance in the field; and Roger Mundry for his precious statistical support. We thank the editor, the associate editor, and two anonymous reviewers for their thoughtful and insightful comments. This research was supported by the Deutsche Forschungsgemeinschaft (DFG, German Research Foundation, Project number 254142454/GRK 2070).

## Supplementary Information for

### Supplementary Information Text

#### Supplementary appendix 1 - age category assessment and criteria for subject selection

We observed all adult and subadult (i.e., small and large subadult) males belonging to the two study parties (party 6 and party 9). Independent observers differentiated developmental stages and assessed age categories using physical markers on a monthly basis (see tables S1 and S2). Males were introduced as focal subjects when they reached subadulthood (∼ 6 years old), as subadult males already establish close affiliations with other males [1] and display male-male strong bonds and coalitionary support [2,3]. Note that, in the light of more accurate aging data, the male age categories previously used for adolescent and adult males [e.g. 3] are updated and improved here. Based on newly acquired data on males that have known birthdates, we now merged the categories for large juvenile and small subadult males into a single category called ‘small subadult males’. This is based on the fact that males in the two previous age categories are already 6 years old, are visibly bigger than adult females, and tend to show testicular enlargement. In addition, this update is in agreement with the criteria used for subadult categories in closely related species (e.g. geladas, [4]; yellow baboons, Amboseli Baboon Research Project Monitoring Guide: https://amboselibaboons.nd.edu/assets/384683/abrp_monitoring_guide_9april2020.pdf).

All males that were associated with the two study parties during the study period and that were present in the study parties for more than 93 days (∼3 month) were included (n=30). Following this rule, two adult males that transferred to and away from party 9 for just over a month (36 presence days) were excluded from all analyses. Additionally, we used the same time criterium within each year to only include males that were present a minimum of 93 days within each yearly dataset. Following this rule, two males that became focal subjects because they reached the ‘small subadult males’ age category at the end of a study year (< 93 presence days), were only included in the following year due to the limited amount of observations in their first year as small subadult males.

In addition to our subject selection criterium, the party composition changed due to demographical changes such as age category changes, party transfer from/to our study parties, and death/disappearance. Overall, seven males that transitioned to small subadult status and two males that transferred into the two study parties were included as focal subjects later in the study, while a total of eight males disappeared during the study period (seven likely due to predation and one due to migration to another gang). See figure S1 for a visual representation of male presence and age category changes during the study period.

#### Supplementary appendix 2 - Genetic analysis: number and type of analysed samples

When DNA extracts from tissue samples were available from previous studies (n=17; see [5]), at least one additional fecal sample was fully genotyped in order to cross-check individual identity. To rule out identification errors during fecal sample collection, for all remaining individuals (n=101) up to five samples were extracted and genotyped (mean=2.86, range=1 to 5). While for 64 individuals, available extracts were fully genotyped and compared for mismatches, for the remaining 37 individuals we fully genotyped only the best quality extract and test-genotyped the remaining extracts at 6 microsatellite loci (i.e. we checked for mismatches across extracts in 6 loci and considered the extract to belong to the same individual only if no mismatches were present; also see Stadele et al. [6] for a similar methodology). After extract exclusion due to mismatches (8 on 289 extracts excluded), the average number of available samples per individual was 2.78 (range=1 to 4). As the offspring sampling is more difficult and therefore more likely to include a greater number of identification errors compared to the sampling for adolescents/adults, all offspring samples were additionally validated using a PCR-based sexing assay to confirm the reported sex (see supplementary appendix 3). The results were in agreement with the reported sex for all samples.

#### Supplementary appendix 3 - Genetic analysis: sex-determination protocol

For sexing, two primers were used to amplify a region of Dead Box gene (F: GGA CGR ACT CTA GAT CGG, R: GTN CAG ATC TAR GAG GAA). The primers amplify one fragment in the female and two fragments in the male. Sexing-PCR was carried out in a 20µl volume using the QIAGEN Multiplex PCR Kit. Reactions contained 10µl of 2x QIAGEN Multiplex PCR Master Mix, 0.15µl BT, 1µl of each primer (10pM) and 0,5-5µl ONA. Water was added as needed to reach the final volume. PCR conditions comprised a pre-denaturing and polymerase activation step at 94°C for 15min, followed by 40-50 cycles at 94°C for 30sec, annealing at 58°C for 30sec and 72°C for 30sec. A final extension step was carried out at 72°C for 5 minutes. Negative controls (only water added) and positive controls (high quality DNA of known male and female sex) were carried along for all amplifications. Sex determination was done by visual inspection of the PCR products on a 2.5% Agarose gel stained with ethidium bromide.

#### Supplementary appendix 4 - Data analysis and modeling supplementary information

Before each analysis, we assessed the number of available data points per estimated term (including the intercept, all random effects, and the residual standard deviation). Note that this estimate can be considered conservative as it includes all estimated terms. We determined model complexity to be adequate for all four models with ratios (data points per term) ranging from 10.11 to 56.40.

To reduce type I error rates, we initially used the maximal random-effect structure comprising all theoretically identifiable random slope components. These also included parameters for the correlations between random intercepts and slopes [7,8]. In all models, however, such correlations were “unidentifiable” (i.e. absolute correlation parameter ∼1; [9]) and were therefore excluded from our final models. Comparisons of log-likelihoods values suggest that model fit was only minimally affected in all cases (tables S5-S10). To ease model conversion and estimate comparison, all covariates were z-transformed to a mean of zero and a standard deviation of one prior to fitting each model [10]. Detailed information about sample size, data standardization/transformation (including means and standard deviations of original values), and the use of non-default optimizers can be found in the supplementary model tables (tables S5-S10).

As our datasets included many zero values in the response variable, for all Poisson models we checked the potential need of zero-inflation by sampling 1000 times as many fitted values as the number of residuals of our models and comparing the number of zeros present in each sample with the number of zero present in our original datasets. We additionally used the function ‘check_zeroinflation’ of the ‘performance’ R package (version 0.7.1; [11]) to check if the ratio of observed and predicted zeros is within the tolerance range. Both methods showed that all models were not underfitting zeros and therefore no zero-inflation was required.

We assessed model stability using a function kindly provided by Roger Mundry, which compares estimates obtained by running models excluding the levels of the random effects one at a time [12]. All models displayed good to adequate levels of stability.

**Fig. S1.**
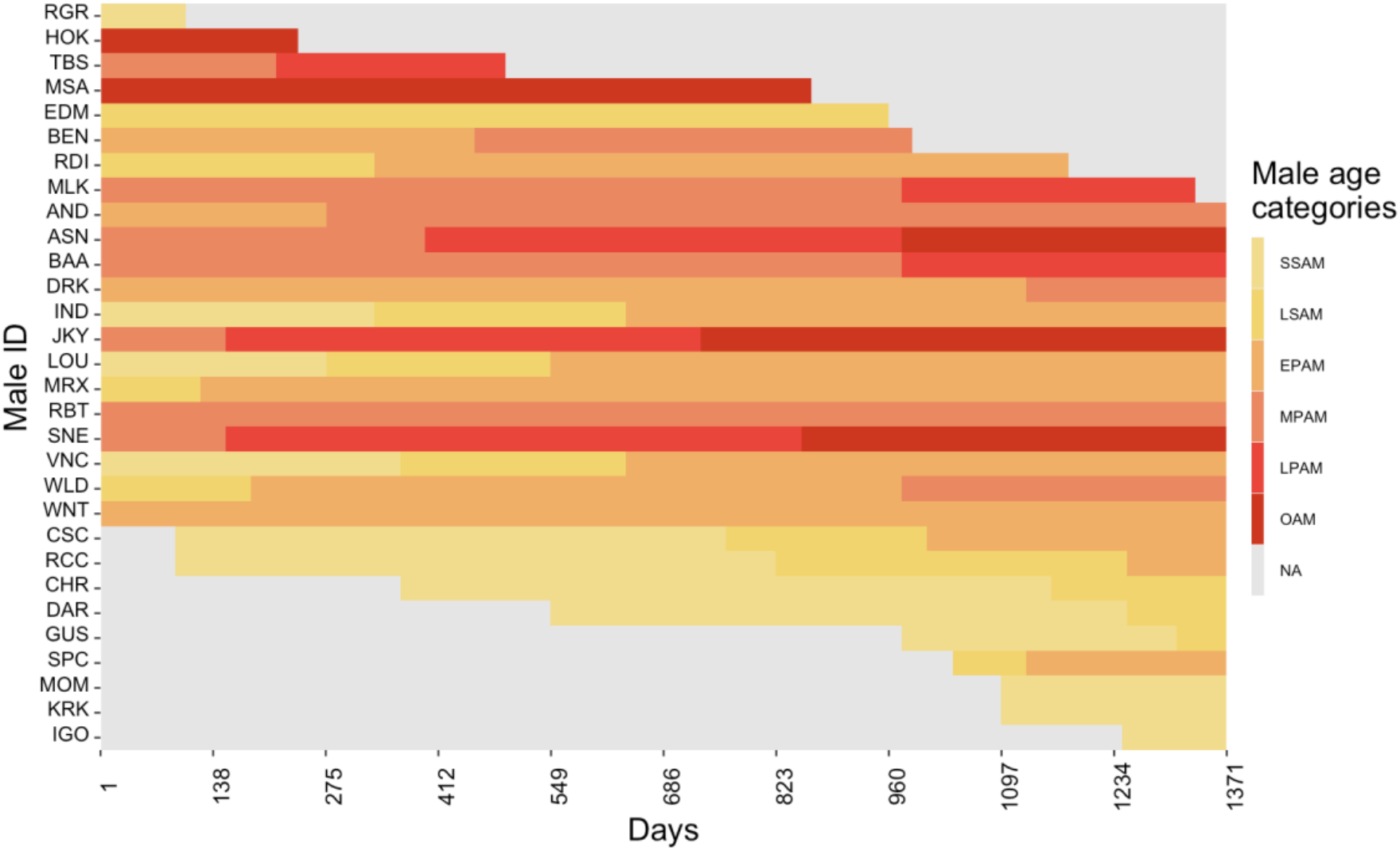
Visual representation of male age categories and male presence for the 30 study subjects during the study period (April 2014 - December 2017). NA (not assessed - in grey) indicates days when males were not present due to demographical changes (i.e. not associated to the study parties, not in the selected age categories, or deceased, also see supplementary appendix 1). Age categories are the following: SSAM= small subadult male; LSAM= large subadult male; EPAM= early prime adult male; MPAM= middle prime adult male; LPAM= late prime adult male; OAM= old adult male.

**Table S1.** Age category definitions for adolescent and adult males.

|  |  |  |
| --- | --- | --- |
| **Adolescent** | SSAM<br>“Small subadult male” | Individuals are visibly bigger than adult females, but still have a lanky appearance and are visibly smaller than adult males (i.e. have not yet attained full body size). Males start experiencing testicular enlargement and have an enlarged scrotum by the end of this phase. Canines start extending beyond tooth row and the mantle start to develop marks. Secondary sexual characteristics are partially but not fully developed (mantle, canine ridges, long canine teeth). |
|  | LSAM<br>“Large subadult male” | Individuals no longer have a lanky appearance but have not yet attained full body size or muscle mass. Body shape is more similar to the one of adult than small subadult males. |
| **Adult** | EPAM<br>“Early-prime adult male” | Secondary sexual characteristics and body size are fully developed. The coat is long and shiny. The ischial callosities become square and wide and the butt may take on reddish color. Teeth in category 5. |
|  | MPAM<br>“Mid-prime adult male” | The mantle may show some breaks. The teeth start decaying (categories 3 or 4). |
|  | LPAM<br>“Late-prime adult male” | The mantle starts to thin out. The male has less muscle mass. Teeth in categories 2 or 3. |
|  | OAM<br>“Old adult male” | The mantle thins out visibly. The male has lost most of his muscle mass. Teeth in categories 1 or 2. |

**Table S2.** Tooth status category definitions [from 13].

| Tooth Category | Definition |
| --- | --- |
| 5 | White teeth with sharp unchipped points. |
| 4 | White teeth or slight yellowing on one or two teeth, some chipping or wear on one tooth. |
| 3 | Some discoloration on several teeth, breaks chipping or tooth wear very evident. |
| 2 | Extensive discoloration, one or both canines missing or broken. |
| 1 | Extensive discoloration, one or both canines missing or worn to the level of premolars and substantial damage to other teeth. |

**Table S3.**
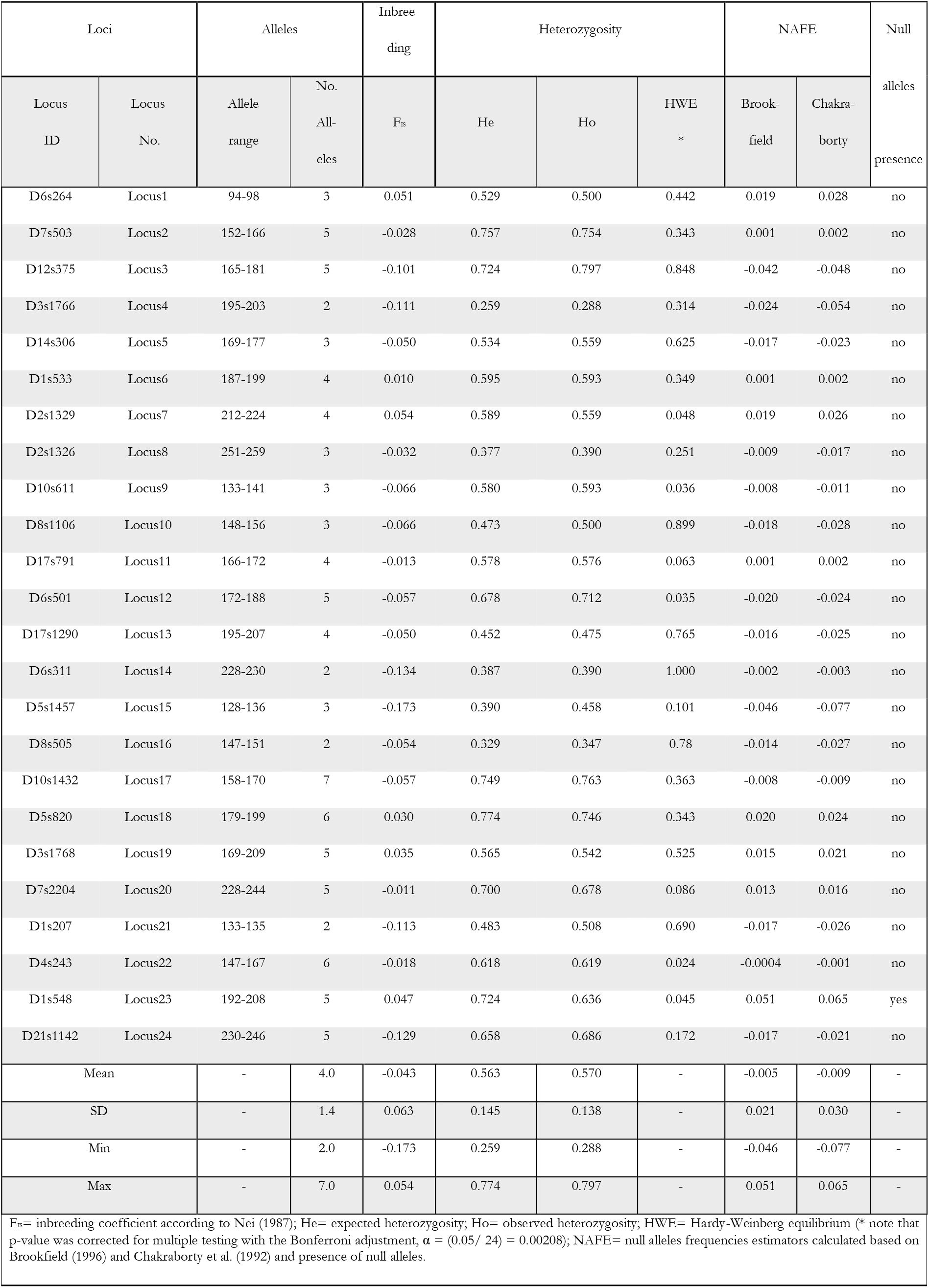
Characteristics of the 24 microsatellite loci used to estimate paternity. (calculated using the genotypes of all males, females and offspring included in the analysis; n=118).

**Table S4.**
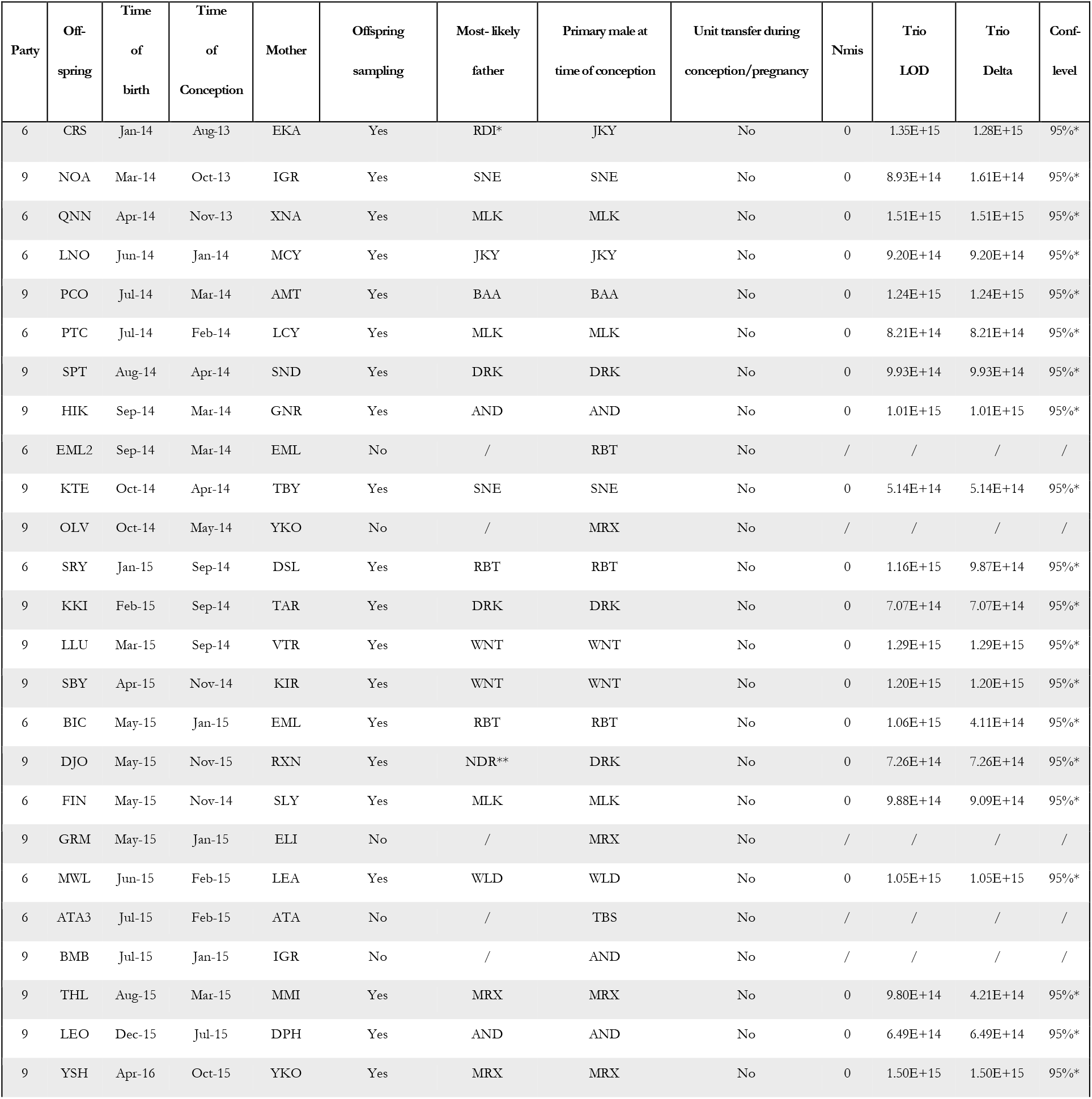
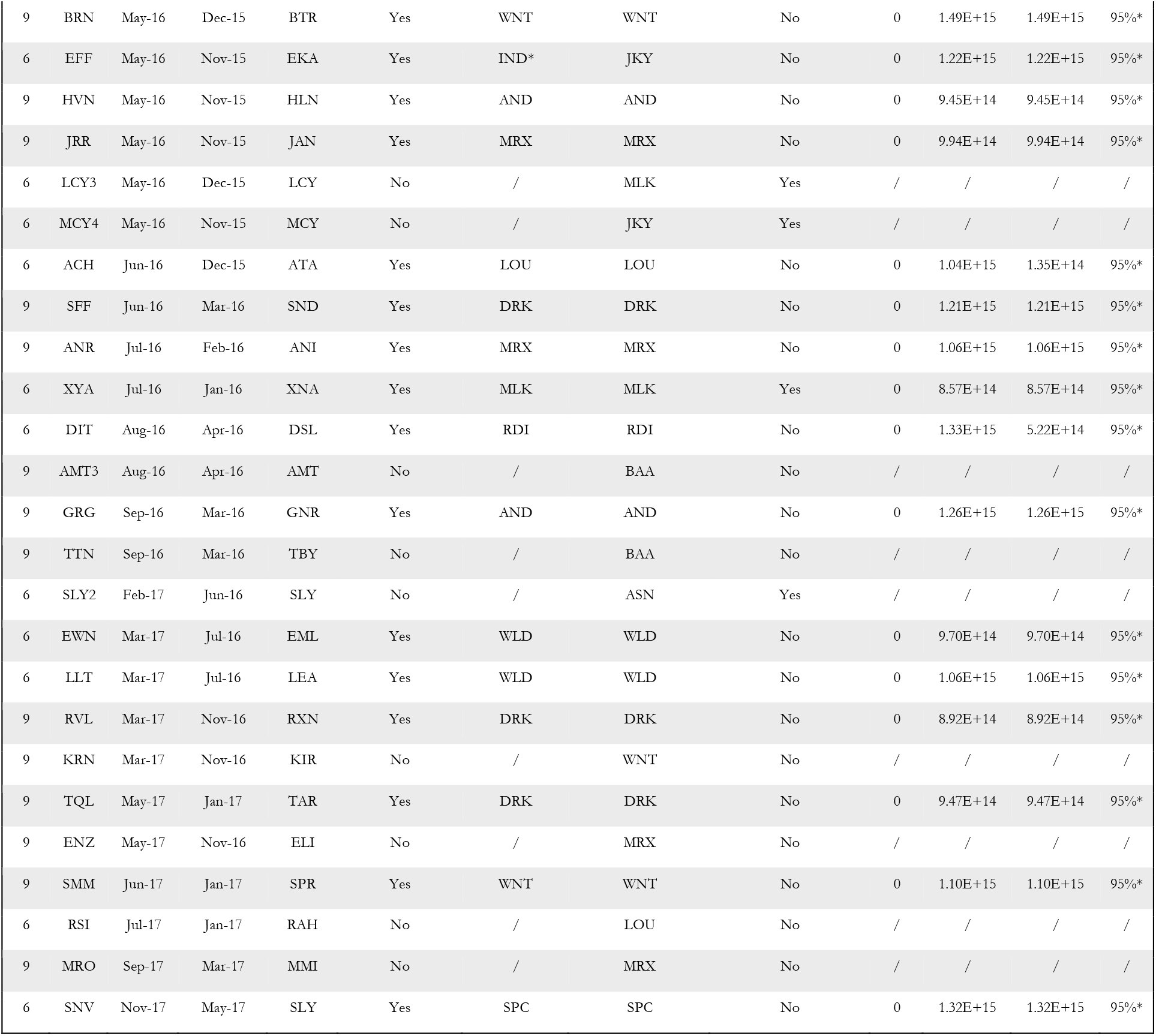
Results of the paternity analysis for offspring born during the study period (2014-2017) in party 6 and 9. Samples and genotypes were available for a total of 36 offspring, while for the other 14 offspring no genetic information were available (see the column ‘Offspring sampling’). Results from the paternity analysis conducted with Cervus 3.0 (version 3.0.7; [14]). The offspring, mother and most likely father identity are reported in the table per each study party. Date (month and year) of conception, date of birth, and identity of the primary male at the time of conception are also reported. Transfer of mothers to another primary male’s unit between time of conception and birth occurred in only 16.7% of cases (see the column ‘Unit transfer during conception/pregnancy’). Nmis indicates number of mismatches; Trio LOD indicates the scores of the logarithm of the likelihood ratio; trio Delta is defined as the difference in LOD scores between the most likely and the second most likely candidate father. The confidence level of the Cervus paternity assignments was set to 95% (‘strict’ criterion). An asterisk in the confidence level column indicates a statistical confidence on paternity assignment higher than 95%. An asterisk in the most likely father column indicates fathers that were not the primary male at the time of conception. In particular, one asterisk indicate a father belonging to the same party and two asterisks a father belonging to a different party of the same gang.

| Party | Offspring | Time of birth | Time of Conception | Mother | Offspring sampling | Most-likely father | Primary male at time of conception | Unit transfer during conception/pregnancy | Nmis | Trio LOD | Trio Delta | Conf-level |
| --- | --- | --- | --- | --- | --- | --- | --- | --- | --- | --- | --- | --- |
| 6 | CRS | Jan-14 | Aug-13 | EKA | Yes | RDI* | JKY | No | 0 | 1.35E+15 | 1.28E+15 | 95%* |
| 9 | NOA | Mar-14 | Oct-13 | IGR | Yes | SNE | SNE | No | 0 | 8.93E+14 | 1.61E+14 | 95%* |
| 6 | QNN | Apr-14 | Nov-13 | XNA | Yes | MLK | MLK | No | 0 | 1.51E+15 | 1.51E+15 | 95%* |
| 6 | LNO | Jun-14 | Jan-14 | MCY | Yes | JKY | JKY | No | 0 | 9.20E+14 | 9.20E+14 | 95%* |
| 9 | PCO | Jul-14 | Mar-14 | AMT | Yes | BAA | BAA | No | 0 | 1.24E+15 | 1.24E+15 | 95%* |
| 6 | PTC | Jul-14 | Feb-14 | LCY | Yes | MLK | MLK | No | 0 | 8.21E+14 | 8.21E+14 | 95%* |
| 9 | SPT | Aug-14 | Apr-14 | SND | Yes | DRK | DRK | No | 0 | 9.93E+14 | 9.93E+14 | 95%* |
| 9 | HIK | Sep-14 | Mar-14 | GNR | Yes | AND | AND | No | 0 | 1.01E+15 | 1.01E+15 | 95%* |
| 6 | EML2 | Sep-14 | Mar-14 | EML | No | / | RBT | No | / | / | / | / |
| 9 | KTE | Oct-14 | Apr-14 | TBY | Yes | SNE | SNE | No | 0 | 5.14E+14 | 5.14E+14 | 95%* |
| 9 | OLV | Oct-14 | May-14 | YKO | No | / | MRX | No | / | / | / | / |
| 6 | SRY | Jan-15 | Sep-14 | DSL | Yes | RBT | RBT | No | 0 | 1.16E+15 | 9.87E+14 | 95%* |
| 9 | KKI | Feb-15 | Sep-14 | TAR | Yes | DRK | DRK | No | 0 | 7.07E+14 | 7.07E+14 | 95%* |
| 9 | LLU | Mar-15 | Sep-14 | VTR | Yes | WNT | WNT | No | 0 | 1.29E+15 | 1.29E+15 | 95%* |
| 9 | SBY | Apr-15 | Nov-14 | KIR | Yes | WNT | WNT | No | 0 | 1.20E+15 | 1.20E+15 | 95%* |
| 6 | BIC | May-15 | Jan-15 | EML | Yes | RBT | RBT | No | 0 | 1.06E+15 | 4.11E+14 | 95%* |
| 9 | DJO | May-15 | Nov-15 | RXN | Yes | NDR** | DRK | No | 0 | 7.26E+14 | 7.26E+14 | 95%* |
| 6 | FIN | May-15 | Nov-14 | SLY | Yes | MLK | MLK | No | 0 | 9.88E+14 | 9.09E+14 | 95%* |
| 9 | GRM | May-15 | Jan-15 | ELI | No | / | MRX | No | / | / | / | / |
| 6 | MWL | Jun-15 | Feb-15 | LEA | Yes | WLD | WLD | No | 0 | 1.05E+15 | 1.05E+15 | 95%* |
| 6 | ATA3 | Jul-15 | Feb-15 | ATA | No | / | TBS | No | / | / | / | / |
| 9 | BMB | Jul-15 | Jan-15 | IGR | No | / | AND | No | / | / | / | / |
| 9 | THL | Aug-15 | Mar-15 | MMI | Yes | MRX | MRX | No | 0 | 9.80E+14 | 4.21E+14 | 95%* |
| 9 | LEO | Dec-15 | Jul-15 | DPH | Yes | AND | AND | No | 0 | 6.49E+14 | 6.49E+14 | 95%* |
| 9 | YSH | Apr-16 | Oct-15 | YKO | Yes | MRX | MRX | No | 0 | 1.50E+15 | 1.50E+15 | 95%* |
| 9 | BRN | May-16 | Dec-15 | BTR | Yes | WNT | WNT | No | 0 | 1.49E+15 | 1.49E+15 | 95%* |
| 6 | EFF | May-16 | Nov-15 | EKA | Yes | IND* | JKY | No | 0 | 1.22E+15 | 1.22E+15 | 95%* |
| 9 | HVN | May-16 | Nov-15 | HLN | Yes | AND | AND | No | 0 | 9.45E+14 | 9.45E+14 | 95%* |
| 9 | JRR | May-16 | Nov-15 | JAN | Yes | MRX | MRX | No | 0 | 9.94E+14 | 9.94E+14 | 95%* |
| 6 | LCY3 | May-16 | Dec-15 | LCY | No | / | MLK | Yes | / | / | / | / |
| 6 | MCY4 | May-16 | Nov-15 | MCY | No | / | JKY | Yes | / | / | / | / |
| 6 | ACH | Jun-16 | Dec-15 | ATA | Yes | LOU | LOU | No | 0 | 1.04E+15 | 1.35E+14 | 95%* |
| 9 | SFF | Jun-16 | Mar-16 | SND | Yes | DRK | DRK | No | 0 | 1.21E+15 | 1.21E+15 | 95%* |
| 9 | ANR | Jul-16 | Feb-16 | ANI | Yes | MRX | MRX | No | 0 | 1.06E+15 | 1.06E+15 | 95%* |
| 6 | XYA | Jul-16 | Jan-16 | XNA | Yes | MLK | MLK | Yes | 0 | 8.57E+14 | 8.57E+14 | 95%* |
| 6 | DIT | Aug-16 | Apr-16 | DSL | Yes | RDI | RDI | No | 0 | 1.33E+15 | 5.22E+14 | 95%* |
| 9 | AMT3 | Aug-16 | Apr-16 | AMT | No | / | BAA | No | / | / | / | / |
| 9 | GRG | Sep-16 | Mar-16 | GNR | Yes | AND | AND | No | 0 | 1.26E+15 | 1.26E+15 | 95%* |
| 9 | TTN | Sep-16 | Mar-16 | TBY | No | / | BAA | No | / | / | / | / |
| 6 | SLY2 | Feb-17 | Jun-16 | SLY | No | / | ASN | Yes | / | / | / | / |
| 6 | EWN | Mar-17 | Jul-16 | EML | Yes | WLD | WLD | No | 0 | 9.70E+14 | 9.70E+14 | 95%* |
| 6 | LLT | Mar-17 | Jul-16 | LEA | Yes | WLD | WLD | No | 0 | 1.06E+15 | 1.06E+15 | 95%* |
| 9 | RVL | Mar-17 | Nov-16 | RXN | Yes | DRK | DRK | No | 0 | 8.92E+14 | 8.92E+14 | 95%* |
| 9 | KRN | Mar-17 | Nov-16 | KIR | No | / | WNT | No | / | / | / | / |
| 9 | TQL | May-17 | Jan-17 | TAR | Yes | DRK | DRK | No | 0 | 9.47E+14 | 9.47E+14 | 95%* |
| 9 | ENZ | May-17 | Nov-16 | ELI | No | / | MRX | No | / | / | / | / |
| 9 | SMM | Jun-17 | Jan-17 | SPR | Yes | WNT | WNT | No | 0 | 1.10E+15 | 1.10E+15 | 95%* |
| 6 | RSI | Jul-17 | Jan-17 | RAH | No | / | LOU | No | / | / | / | / |
| 9 | MRO | Sep-17 | Mar-17 | MMI | No | / | MRX | No | / | / | / | / |
| 6 | SNV | Nov-17 | May-17 | SLY | Yes | SPC | SPC | No | 0 | 1.32E+15 | 1.32E+15 | 95%* |

**Table S5.** Model 1 - Male-male dyadic composite sociality index (DSI) and coalitionary support. Model table of male-male OSI’s effect on coalitionary rate per contact hour.

| Model formula:<br>Glmer(Coalition_Count ~ z.log.DSI + Party + Year +<br>(1 + z.log.DSI + Year_code.2 + Year_code.3 + Year_code.4 \|\| Subject_ID) + (1 + z.log.DSI + Year_code.2 + Year_code.3 + Year_code.4 \|\| Partner_ID) + (1 \|\| Dyad_Subject+Partner_ID) + offset(log(Dyadic_Contact_Hours)), family= “poisson”, data=d, control=Contr)<br>Contr=glmerControl(optimizer=“bobyqa”, optCtrl=list(maxfun=100000)) |  |  |  |  |  |  |  |
| --- | --- | --- | --- | --- | --- | --- | --- |
| | Estimate | Std. Error | 2.5% CI | 97.5 % CI | $\chi^2$ | df | Pr(Chi) |
| Intercept | -9.429 | 0.375 | -10.495 | -8.857 | (3) | (3) | (3) |
| z.log.DSI<sup>(1)</sup> | 0.781 | 0.108 | 0.500 | 0.994 | 27.760 | 1 | <0.001 |
| Party 9<sup>(2)</sup> | 0.332 | 0.346 | -0.341 | 0.993 | 0.919 | 1 | 0.338 |
| Year_2015<sup>(2)</sup> | 0.430 | 0.375 | -0.425 | 1.205 |  |  |  |
| Year_2016<sup>(2)</sup> | -0.588 | 0.524 | -1.756 | 0.392 | 14.174 | 3 | 0.003 |
| Year_2017<sup>(2)</sup> | -1.520 | 0.421 | -2.475 | -0.638 |  |  |  |
| Estimates calculated from the generalized linear mixed model with standard errors. Models examine the effects or predictors indicated on the left on coalitionary rate per contact hour (main predictors above the dotted line; control predictors below). CI = confidence interval.<br><u>The sample for this model consisted of:</u> number of obs: 958; groups: Dyad_Subject+Partner_ID=388; Subject_ID=30; Partner_ID=30.<br><u>Dispersion parameter</u> =0.289 (no concern for overdispersion; value < 1.0).<br>Effect sizes - R<sup>2</sup>: R2m= 0.155; R2c= 0.706; calculated with the method “Trigamma” and the “r.squaredGLMM” function of the “MuMIn” R package<br>Comparisons of log-likelihoods between model with/without correlations between random intercepts and slopes: full model including the correlation parameters: -449.4 (df=37); full model lacking the correlation parameters: -459.8215 (df=17)<br>(1) The predictor of interest DSI was log-transformed and afterwards z-transformed to a mean of 0 and a standard deviation (SD) of 1; original mean (SD): 1.000 (2.291); log-transformed mean (SD): 0.399 (0.641).<br>(2) Year was dummy coded with 2014 being the reference category. Party was dummy coded with Party 6 being the reference category.<br>(3) Not shown due to very limited interpretability. |  |  |  |  |  |  |  |

**Table S6.** Model 2 - Mode number of associated females and male-male sociality (male bond strength and number of strong bonds). Model table of male bond strength and number of strong bonds effect on the mode number of associated females (calculated as mode per male per year).

| Model formula: |  |  |  |  |  |  |  |
| --- | --- | --- | --- | --- | --- | --- | --- |
| Glmer(Nfemales.mode ~ z.Bond_Strenght_top3DSI + z.N_StrongBonds + Party + Year + (1 + z.Bond_Strenght_top3DSI \|\| Male_ID), family= "poisson", data=d, control=Contr) |  |  |  |  |  |  |  |
| Contr=glmerControl(optimizer="bobyqa", optCtrl=list(maxfun=100000)) |  |  |  |  |  |  |  |
| | Estimate | Std. Error | 2.5% CI | 97.5 % CI | $\chi^2$ | df | Pr(Chi) |
| Intercept | -0.625 | 0.372 | -1.522 | 0.017 | (3) | (3) | (3) |
| z.Bond_Strenght_top3DSI<sup>(1)</sup> | -0.749 | 0.266 | -1.381 | -0.254 | 8.985 | 1 | 0.003 |
| z.N_StrongBonds<sup>(1)</sup> | -0.288 | 0.220 | -0.779 | 0.176 | 1.793 | 1 | 0.181 |
| Party 9<sup>(2)</sup> | 0.191 | 0.446 | -0.665 | 1.040 | 0.177 | 1 | 0.674 |
| Year_2015<sup>(2)</sup> | -0.100 | 0.290 | -0.791 | 0.498 |  |  |  |
| Year_2016<sup>(2)</sup> | -0.263 | 0.301 | -1.000 | 0.321 | 1.319 | 3 | 0.725 |
| Year_2017<sup>(2)</sup> | -0.320 | 0.308 | -0.952 | 0.267 |  |  |  |
| Estimates calculated from the generalized linear mixed model with standard errors. Models examine the effects of predictors indicated on the left on mode number of associated females (main predictors above the dotted line; control predictors below). CI = confidence interval. |  |  |  |  |  |  |  |
| <u>The sample for this model consisted of:</u> number of obs: 91; groups: Male_ID=30. |  |  |  |  |  |  |  |
| <u>Dispersion parameter</u> = 0.542 (no concern for overdispersion; value < 1.0). |  |  |  |  |  |  |  |
| Effect sizes - R<sup>2</sup>: R2m= 0.335; R2c= 0.593; calculated with the method "Trigamma" and the "r.squaredGLMM" function of the "MuMIn" R package |  |  |  |  |  |  |  |
| Comparisons of log-likelihoods between model with/without correlations between random intercepts and slopes: full model including the correlation parameter: -101.436 (df=10); full model lacking the correlation parameter: -103.471 (df=9) |  |  |  |  |  |  |  |
| (1) The predictors of interest were z-transformed to a mean of 0 and a standard deviation (SD) of 1; original mean (SD): Bond_Strenght_top3DSI 9.353 (6.506); N_StrongBonds 2.176 (1.525). |  |  |  |  |  |  |  |
| (2) Year was dummy coded with 2014 being the reference category. Party was dummy coded with Party 6 being the reference category. |  |  |  |  |  |  |  |
| (3) Not shown due to very limited interpretability. |  |  |  |  |  |  |  |

**Table S7.** Model 3 - Number of sired offspring and male-male sociality (male bond strength and number of strong bonds). Model table of male bond strength and number of strong bonds effect on number of sired offspring (calculated as count per male per year).

| Model formula: |  |  |  |  |  |  |  |
| --- | --- | --- | --- | --- | --- | --- | --- |
| Glmer(NsiredOffspring ~ z.Bond_Strenght_top3DSI + z.N_StrongBonds + Party + Year +<br>(1 + z.Bond_Strenght_top3DSI \|\| Male_ID), family= "poisson", data=d, control=Contr)<br>Contr=glmerControl(optimizer="bobyqa", optCtrl=list(maxfun=100000)) |  |  |  |  |  |  |  |
| | Estimate | Std.<br>Error | 2.5% CI | 97.5 % CI | $\chi^2$ | df | Pr(Chi) |
| Intercept | -1.035 | 0.349 | -1.969 | -0.495 | (3) | (3) | (3) |
| z.Bond_Strenght_top3DSI<sup>(1)</sup> | -0.655 | 0.288 | -1.358 | -0.144 | 5.738 | 1 | 0.017 |
| z.N_StrongBonds<sup>(1)</sup> | -0.085 | 0.245 | -0.629 | 0.367 | 0.122 | 1 | 0.727 |
| Party 9<sup>(2)</sup> | 0.288 | 0.295 | -0.315 | 0.959 | 0.936 | 1 | 0.333 |
| Year_2015<sup>(2)</sup> | 0.078 | 0.418 | -0.796 | 1.005 |  |  |  |
| Year_2016<sup>(2)</sup> | 0.260 | 0.399 | -0.553 | 1.146 | 1.205 | 3 | 0.752 |
| Year_2017<sup>(2)</sup> | -0.168 | 0.429 | -1.140 | 0.806 |  |  |  |
| Estimates calculated from the generalized linear mixed model with standard errors. Models examine the effects of predictors indicated on the left on number of sired offspring (main predictors above the dotted line; control predictors below). CI = confidence interval. |  |  |  |  |  |  |  |
| <u>The sample for this model consisted of:</u> number of obs: 91; groups: Male_ID=30. |  |  |  |  |  |  |  |
| <u>Dispersion parameter</u> = 1.159 (no substantial concern for overdispersion; value ~1.0) |  |  |  |  |  |  |  |
| Effect sizes - R<sup>2</sup>: R2m= 0.106; R2c= 0.106; calculated with the method "Trigamma" and the "r.squaredGLMM" function of the "MuMIn" R package |  |  |  |  |  |  |  |
| Comparisons of log-likelihoods between model with/without correlations between random intercepts and slopes: full model including the correlation parameter: -79.107 (df=10); full model lacking the correlation parameter: -79.672 (df=9) |  |  |  |  |  |  |  |
| (1) The predictors of interest were z-transformed to a mean of 0 and a standard deviation (SD) of 1; original mean (SD): Bond_Strenght_top3DSI 9.353 (6.506); N_StrongBonds 2.176 (1.525). |  |  |  |  |  |  |  |
| (2) Year was dummy coded with 2014 being the reference category. Party was dummy coded with Party 6 being the reference category. |  |  |  |  |  |  |  |
| (3) Not shown due to very limited interpretability. |  |  |  |  |  |  |  |

**Table S8.** Model 4 - Post-hoc test: time males spent affiliating with other males by number of associated females. Model table of male number of associated females’ effect on proportion of time males spent affiliating with other males (i.e. grooming plus contact-sit).

| Model formula: |  |  |  |  |  |  |
| --- | --- | --- | --- | --- | --- | --- |
| glmmTMB(tr.prop.aff ~ z.FocalMaleNfemales + Party + Year + (1\|Male_ID), family = beta_family(link="logit"), data=d) |  |  |  |  |  |  |
|  | Estimate | Std. Error | 2.5% CI | 97.5 % CI | z-value | p-value |
| Intercept | -3.645 | 0.22 | -4.04 | -3.277 | (3) | (3) |
| z.FocalMaleNfemales<sup>(1)</sup> | -0.371 | 0.107 | -0.55 | -0.209 | -3.479 | 0.001 |
| Party 9<sup>(2)</sup> | -0.089 | 0.212 | -0.442 | 0.289 | -0.418 | 0.676 |
| Year_2015<sup>(2)</sup> | 0.399 | 0.209 | 0.032 | 0.758 | 1.908 | 0.056 |
| Year_2016<sup>(2)</sup> | 0.237 | 0.227 | -0.15 | 0.644 | 1.046 | 0.296 |
| Year_2017<sup>(2)</sup> | 0.186 | 0.223 | -0.196 | 0.566 | 0.836 | 0.403 |
| Estimates calculated from the generalized linear mixed model with standard errors. Models examine the effects of predictors indicated on the left on proportion of time males spent affiliating with other males (main predictors above the dotted line; control predictors below). CI = confidence interval. |  |  |  |  |  |  |
| <u>The sample for this model consisted of:</u> number of obs: 147; groups: Male_ID=30. |  |  |  |  |  |  |
| <u>Dispersion parameter</u> = 1.283 (Moderate overdispersion) - Std. Error, Z- and p-values were corrected by the overdispersion level according to Gelman and Hill (2007) |  |  |  |  |  |  |
| Effect sizes - R<sup>2</sup>: R2m= 0.197; R2c= 0.290; calculated with the "r2" function of the "performance" R package |  |  |  |  |  |  |
| (1) The predictor of interest FocalMaleNfemales was z-transformed to a mean of 0 and a standard deviation (SD) of 1; original mean (SD): 1.442 (1.513). (2) Year was dummy coded with 2014 being the reference category. Party was dummy coded with Party 6 being the reference category. |  |  |  |  |  |  |
| (3) Not shown due to very limited interpretability. |  |  |  |  |  |  |

**Table S9.** Subset of Model 2 - Mode number of associated females and male-male sociality (male bond strength and number of strong bonds). This model was ran with a subset of data only including large subadult and adult males within each year - Model table of male bond strength and number of strong bonds effect on the mode number of associated females (calculated as mode per male per year).

| Model formula: |  |  |  |  |  |  |  |
| --- | --- | --- | --- | --- | --- | --- | --- |
| Glmer(Nfemales.mode ~ z.Bond_Strenght_top3DSI + z.N_StrongBonds + Party + Year +<br>(1 + z.Bond_Strenght_top3DSI \|\| Male_ID), family= "poisson", data=Subset_d, control=Contr)<br>Contr=glmerControl(optimizer="bobyqa", optCtrl=list(maxfun=100000)) |  |  |  |  |  |  |  |
| | Estimate | Std.<br>Error | 2.5% CI | 97.5 % CI | $\chi^2$ | df | Pr(Chi) |
| Intercept | -0.229 | 0.321 | -0.969 | 0.379 | (3) | (3) | (3) |
| z.Bond_Strenght_top3DSI<sup>(1)</sup> | -0.739 | 0.259 | -1.368 | -0.281 | 9.246 | 1 | 0.002 |
| z.N_StrongBonds<sup>(1)</sup> | -0.201 | 0.214 | -0.676 | 0.217 | 0.921 | 1 | 0.337 |
| Party 9<sup>(2)</sup> | 0.294 | 0.365 | -0.431 | 1.036 | 0.602 | 1 | 0.438 |
| Year_2015<sup>(2)</sup> | -0.132 | 0.287 | -0.787 | 0.431 |  |  |  |
| Year_2016<sup>(2)</sup> | -0.285 | 0.297 | -0.933 | 0.361 | 1.234 | 3 | 0.745 |
| Year_2017<sup>(2)</sup> | -0.291 | 0.302 | -0.893 | 0.377 |  |  |  |
| Estimates calculated from the generalized linear mixed model with standard errors. Models examine the effects of predictors indicated on the left on mode number of associated females (main predictors above the dotted line; control predictors below). CI = confidence interval. |  |  |  |  |  |  |  |
| <u>The sample for this model consisted of:</u> number of obs: 76; groups: Male_ID=25. |  |  |  |  |  |  |  |
| <u>Dispersion parameter</u> = 0.628 (no concern for overdispersion; value < 1.0). |  |  |  |  |  |  |  |
| Effect sizes - R<sup>2</sup>: R2m= 0.382; R2c= 0.540; calculated with the method "Trigamma" and the "r.squaredGLMM" function of the "MuMIn" R package |  |  |  |  |  |  |  |
| Comparisons of log-likelihoods between model with/without correlations between random intercepts and slopes: full model including the correlation parameter: -95.465 (df=10); full model lacking the correlation parameter: -96.274 (df=9) |  |  |  |  |  |  |  |
| (1) The predictors of interest were z-transformed to a mean of 0 and a standard deviation (SD) of 1; original mean (SD): Bond_Strenght_top3DSI 9.190 (7.038); N_StrongBonds 2.053 (1.582). |  |  |  |  |  |  |  |
| (2) Year was dummy coded with 2014 being the reference category. Party was dummy coded with Party 6 being the reference category. |  |  |  |  |  |  |  |
| (3) Not shown due to very limited interpretability. |  |  |  |  |  |  |  |

**Table S10.** Subset of Model 3 - Number of sired offspring and male-male sociality (male bond strength and number of strong bonds). This model was ran with a subset of data only including large subadult and adult males within each year - Model table of male bond strength and number of strong bonds effect on number of sired offspring (calculated as count per male per year).

| Model formula: |  |  |  |  |  |  |  |
| --- | --- | --- | --- | --- | --- | --- | --- |
| Glmer(NsiredOffspring ~ z.Bond_Strenght_top3DSI + z.N_StrongBonds + Party + Year +<br>(1 + z.Bond_Strenght_top3DSI \|\| Male_ID), family= "poisson", data=d, control=Contr)<br>Contr=glmerControl(optimizer="bobyqa", optCtrl=list(maxfun=100000)) |  |  |  |  |  |  |  |
| | Estimate | Std.<br>Error | 2.5% CI | 97.5 % CI | $\chi^2$ | df | Pr(Chi) |
| Intercept | -0.761 | 0.351 | -1.705 | -0.195 | (3) | (3) | (3) |
| z.Bond_Strenght_top3DSI<sup>(1)</sup> | -0.664 | 0.289 | -1.372 | -0.151 | 6.197 | 1 | 0.013 |
| z.N_StrongBonds<sup>(1)</sup> | 0.032 | 0.240 | -0.513 | 0.489 | 0.017 | 1 | 0.895 |
| Party 9<sup>(2)</sup> | 0.323 | 0.298 | -0.243 | 1.004 | 1.191 | 1 | 0.275 |
| Year_2015<sup>(2)</sup> | -0.030 | 0.418 | -0.892 | 0.868 |  |  |  |
| Year_2016<sup>(2)</sup> | 0.147 | 0.398 | -0.670 | 1.009 | 0.766 | 3 | 0.858 |
| Year_2017<sup>(2)</sup> | -0.200 | 0.429 | -1.126 | 0.696 |  |  |  |
| Estimates calculated from the generalized linear mixed model with standard errors. Models examine the effects of predictors indicated on the left on number of sired offspring (main predictors above the dotted line; control predictors below). CI = confidence interval. |  |  |  |  |  |  |  |
| <u>The sample for this model consisted of:</u> number of obs: 76; groups: Male_ID=25. |  |  |  |  |  |  |  |
| <u>Dispersion parameter</u> = 1.004 (no substantial concern for overdispersion; value ~1.0) |  |  |  |  |  |  |  |
| Effect sizes - R<sup>2</sup>: R2m= 0.122; R2c= 0.122; calculated with the method "Trigamma" and the "r.squaredGLMM" function of the "MuMIn" R package |  |  |  |  |  |  |  |
| Comparisons of log-likelihoods between model with/without correlations between random intercepts and slopes: full model including the correlation parameter: -73.159 (df=10); full model lacking the correlation parameter: -73.159 (df=9) |  |  |  |  |  |  |  |
| (1) The predictors of interest were z-transformed to a mean of 0 and a standard deviation (SD) of 1; original mean (SD): Bond_Strenght_top3DSI 9.190 (7.038); N_StrongBonds 2.053 (1.582). |  |  |  |  |  |  |  |
| (2) Year was dummy coded with 2014 being the reference category. Party was dummy coded with Party 6 being the reference category. |  |  |  |  |  |  |  |
| (3) Not shown due to very limited interpretability. |  |  |  |  |  |  |  |

## Notes

https://osf.io/7v8r5/?view_only=fbfba4deb6284522b67d955b91d902df

### Competing Interest Statement

The authors have declared no competing interest.

## References

1. Darwin C. 1871 The Descent of Man, and Selection in Relation to Sex. London: Murray.

2. Andersson M. 1994 Sexual Selection. Princeton: Princeton University Press.

3. Clutton-Brock TH. 2016 Mammal Societies. Chichester, West Sussex, UK: John Wiley & Sons.

4. Le Boeuf BJ. 1974 Male-male Competition and Reproductive Success in Elephant Seals. Am Zool 14, 163–176. (doi:10.1093/icb/14.1.163)

5. Paul A. 2002 Sexual selection and mate choice. International Journal of Primatology 23, 877–904. (doi:https://doi.org/10.1023/A:1015533100275)

6. Andersson M, Iwasa Y. 1996 Sexual selection. Trends Ecol Evol. 11, 53–8. (doi:10.1016/0169-5347(96)81042-1)

7. Andersson M. 1982 Female choice selects for extreme tail length in a widowbird. Nature 299, 818–820. (doi:https://doi.org/10.1038/299818a0)

8. Nowicki S, Searcy W. 2004 Song Function and the Evolution of Female Preferences: Why Birds Sing, Why Brains Matter. Ann N Y Acad Sci 1016, 704–23. (doi:10.1196/annals.1298.012)

9. Bygott J, Bertram B, Hanby J. 1979 Male lions in large coalitions gain reproductive advantages. Nature 282, 839–841.

10. Feh C. 1999 Alliances and reproductive success in Camargue stallions. Animal Behaviour 57, 705–713. (doi:https://doi.org/10.1006/anbe.1998.1009)

11. Wiszniewski J, Corrigan S, Beheregaray LB, Möller LM. 2012 Male reproductive success increases with alliance size in Indo-Pacific bottlenose dolphins (Tursiops aduncus). Journal of Animal Ecology 81, 423–431. (doi:10.1111/j.1365-2656.2011.01910.x)

12. Snyder-Mackler N, Alberts SC, Bergman TJ. 2012 Concessions of an alpha male? Cooperative defence and shared reproduction in multi-male primate groups. Proceedings of the Royal Society B: Biological Sciences 279, 3788–3795. (doi:10.1098/rspb.2012.0842)

13. Chowdhury S, Pines M, Saunders J, Swedell L. 2015 The adaptive value of secondary males in the polygynous multi-level society of hamadryas baboons. American Journal of Physical Anthropology 158, 501–513. (doi:10.1002/ajpa.22804)

14. Ryder TB, Parker PG, Blake JG, Loiselle BA. 2009 It takes two to tango: Reproductive skew and social correlates of male mating success in a lek-breeding bird. Proceedings of the Royal Society B: Biological Sciences 276, 2377–2384. (doi:10.1098/rspb.2009.0208)

15. Taborsky M. 1987 Cooperative Behaviour in Fish: Coalitions, Kin Groups and Reciprocity. In Animal Societies: Theories and Facts (eds Y Ito, J Brown, J Kikkawa), pp. 229–237. Tokyo: Japan Sci. Soc. Press.

16. Silk JB. 2002 Using the ‘F’-word in primatology. Behaviour 139, 421–446. (doi:10.1163/156853902760102735)

17. Schülke O, Bhagavatula J, Vigilant L, Ostner J. 2010 Social bonds enhance reproductive success in male macaques. Current Biology 20, 2207–2210. (doi:10.1016/j.cub.2010.10.058)

18. Feldblum JT, Krupenye C, Bray J, Pusey AE, Gilby IC. 2021 Social bonds provide multiple pathways to reproductive success in wild male chimpanzees. iScience 24, 102864. (doi:10.1016/j.isci.2021.102864)

19. Bray J, Feldblum JT, Gilby IC. 2021 Social bonds predict dominance trajectories in adult male chimpanzees. Animal Behaviour 179, 339–354. (doi:10.1016/j.anbehav.2021.06.031)

20. Smuts BB, Smuts RW. 1993 Male-Aggression and Sexual Coercion of Females in Nonhuman-Primates and Other Mammals - Evidence and Theoretical Implications. Advances in the Study of Behavior 22, 1–63.

21. Stanford CB. 1998 Predation and male bonds in primate societies. Behaviour 135, 513–533.

22. Fischer J et al. 2017 Charting the neglected West: The social system of Guinea baboons. American Journal of Physical Anthropology 162, 15–31. (doi:10.1002/ajpa.23144)

23. Patzelt A, Kopp GH, Ndao I, Kalbitzer U, Zinner D, Fischer J. 2014 Male tolerance and male-male bonds in a multilevel primate society. Proceedings of the National Academy of Sciences of the United States of America 111, 14740–14745. (doi:10.1073/pnas.1405811111)

24. Dal Pesco F, Trede F, Zinner D, Fischer J. 2021 Kin bias and male pair-bond status shape male-male relationships in a multilevel primate society. Behav Ecol Sociobiol 75, 24. (doi:10.1007/s00265-020-02960-8)

25. Goffe AS, Zinner D, Fischer J. 2016 Sex and friendship in a multilevel society: behavioural patterns and associations between female and male Guinea baboons. Behavioral Ecology and Sociobiology 70, 323–336. (doi:10.1007/s00265-015-2050-6)

26. Pines M, Saunders J, Swedell L. 2011 Alternative routes to the leader male role in a multi-level society: Follower vs. solitary male strategies and outcomes in hamadryas baboons. American Journal of Primatology 73, 679–691. (doi:10.1002/ajp.20951)

27. Fischer J et al. 2019 Insights into the evolution of social systems and species from baboon studies. eLife 8, e50989. (doi:10.7554/eLife.50989)

28. Dal Pesco F, Fischer J. 2018 Greetings in male Guinea baboons and the function of rituals in complex social groups. Journal of Human Evolution 125, 87–98. (doi:10.1016/j.jhevol.2018.10.007)

29. Dal Pesco F. 2020 Dynamics and fitness benefits of male-male sociality in wild Guinea baboons (Papio papio). PhD Thesis, Georg-August University, Göttingen.

30. Altmann J. 1974 Observational study of behavior: sampling methods. Behaviour 49, 227–266. (doi:10.1163/156853974X00534)

31. Silk JB, Cheney D, Seyfarth R. 2013 A practical guide to the study of social relationships. Evolutionary Anthropology 22, 213–225. (doi:10.1002/evan.21367)

32. Silk JB, Alberts SC, Altmann J. 2003 Social bonds of female baboons enhance infant survival. Science 302, 1231–1234. (doi:10.1126/science.1088580)

33. McFarland R, Murphy D, Lusseau D, Henzi SP, Parker JL, Pollet TV, Barrett L. 2017 The ‘strength of weak ties’ among female baboons: fitness-related benefits of social bonds. Animal Behaviour 126, 101–106. (doi:10.1016/j.anbehav.2017.02.002)

34. Dal Pesco F, Trede F, Zinner D, Fischer J. 2021 Analysis of genetic relatedness and paternity assignment in wild Guinea baboons (Papio papio) based on microsatellites. Version 2. protocols.io (doi:dx.doi.org/10.17504/protocols.io.bsg7nbzn)

35. Kopp GH, Fischer J, Patzelt A, Roos C, Zinner D. 2015 Population genetic insights into the social organization of Guinea baboons (Papio papio): Evidence for female-biased dispersal. American Journal of Primatology 77, 878–889. (doi:10.1002/ajp.22415)

36. Kalinowski ST, Taper ML, Marshall TC. 2007 Revising how the computer program CERVUS accommodates genotyping error increases success in paternity assignment. Molecular Ecology 16, 1099–1106. (doi:10.1111/j.1365-294X.2007.03089.x)

37. R Development Core Team. 2021 R: a language and environment for statistical computing. R Foundation for Statistical Computing Vienna, Austria.

38. RStudio, Inc., Boston M. 2021 RStudio: Integrated Development for R. RStudio, Inc. Boston, Massachusetts.

39. Baayen RH. 2008 Analyzing linguistic data: A practical introduction to statistics using R. Cambridge: Cambridge University Press.

40. Bates D, Mächler M, Bolker B, Walker S. 2015 Fitting Linear Mixed-Effects Models using lme4. Journal of Statistical Software 67, 1–48. (doi:10.18637/jss.v067.i01)

41. Brooks ME, Kristensen K, Koen J, Magnusson A, Berg CW, Nielsen A, Skaug HJ, Maechler M, Bolker BM. 2017 glmmTMB Balances Speed and Flexibility Among Packages for Zero-inflated Generalized Linear Mixed Modeling. The R Journal 9, 378–400. (doi:https://journal.r-project.org/archive/2017/RJ-2017-066/index.html)

42. Barr DJ, Levy R, Scheepers C, Tily HJ. 2013 Random effects structure for confirmatory hypothesis testing: Keep it maximal. Journal of Memory and Language 68, 255–278. (doi:10.1016/j.jml.2012.11.001)

43. Matuschek H, Kliegl R, Vasishth S, Baayen H, Bates D. 2017 Balancing type I error and power in linear mixed models. Journal of Memory and Language 94, 305–315.

44. Schielzeth H. 2010 Simple means to improve the interpretability of regression coefficients. Methods in Ecology and Evolution 1, 103–113. (doi:10.1111/j.2041-210X.2010.00012.x)

45. Field A. 2005 Discovering Statistics using SPSS. London: Sage Publications.

46. Fox J, Weisberg S. 2011 An {R} Companion to Applied Regression. second edition. Thousand Oaks, CA: Sage Publications.

47. Dobson AJ. 2002 An introduction to generalized linear models. 2nd edn. London: Chapman & Hall/CRC.

48. Forstmeier W, Schielzeth H. 2011 Cryptic multiple hypotheses testing in linear models: Overestimated effect sizes and the winner’s curse. Behavioral Ecology and Sociobiology 65, 47–55. (doi:10.1007/s00265-010-1038-5)

49. Barton K. 2020 MuMIn: Multi-Model Inference. R package version 1.43.17.

50. Lüdecke D, Ben-Shachar M, Patil I, Waggoner P, Makowski D. 2021 performance: An R Package for Assessment, Comparison and Testing of Statistical Models. Journal of Open Source Software 6, 3139. (doi:10.21105/joss.03139)

51. McCullagh P, Nelder JA. 1989 Generalized Linear Models. 2nd edn. London: Chapman and Hall.

52. Mielke A, Samuni L. 2021 Accuracy and Precision of Social Relationship Indices. Preprint: http://biorxiv.org/lookup/doi/10.1101/2021.04.25.441321

53. Bolker BM. 2008 Ecological Models and Data in R. Princeton, New Jersey: Princeton University Press.

54. Smithson M, Verkuilen J. 2006 A better lemon squeezer? maximum-likelihood regression with beta-distributed dependent variables. Psychological Methods 11, 54–71.

55. Gelman A, Hill J. 2007 Data Analysis Using Regression and Multilevel/Hierarchical Models. Cambridge, New Jersey: Cambridge University Press.

56. Cameron EZ, Setsaas TH, Linklater WL. 2009 Social bonds between unrelated females increase reproductive success in feral horses. Proceedings of the National Academy of Sciences of the United States of America 106, 13850–13853. (doi:10.1073/pnas.0900639106)

57. Berghänel A, Ostner J, Schröder U, Schülke O. 2011 Social bonds predict future cooperation in male Barbary macaques, Macaca sylvanus. Animal Behaviour 81, 1109–1116. (doi:10.1016/j.anbehav.2011.02.009)

58. Zhu P, Grueter CC, Garber PA, Li D, Xiang Z, Ren B, Li M. 2018 Seasonal changes in social cohesion among males in a same-sex primate group. Am J Primatol e22914.

59. Rathke E-M, Berghaenel A, Bissonnette A, Ostner J, Schülke O. 2017 Age-dependent change of coalitionary strategy in male Barbary macaques. Primate Biology 4, 1–7. (doi:10.5194/pb-4-1-2017)

60. Rosati AG, Hagberg L, Enigk DK, Otali E, Emery Thompson M, Muller MN, Wrangham RW, Machanda ZP. 2020 Social selectivity in aging wild chimpanzees. Science 370, 473–476. (doi:10.1126/science.aaz9129)

61. Kalbitz J, Schülke O, Ostner J. 2017 Triadic male-infant-male interaction serves in bond maintenance in male Assamese macaques. PLoS ONE 12, 16–19. (doi:10.1371/journal.pone.0183981)

62. Gerber L et al. 2019 Affiliation history and age similarity predict alliance formation in adult male bottlenose dolphins. Behavioral Ecology 31, 361–370. (doi:10.1093/beheco/arz195)

63. Kalbitzer U. 2014 Foundations of variation in male aggressiveness and tolerance between chacma baboons (Papio ursinus) in Botswana and Guinea baboons (P. papio) in Senegal. PhD Thesis, Georg-August-Universität Göttingen.

64. Jolly CJ. 2020 Philopatry at the frontier: A demographically driven scenario for the evolution of multilevel societies in baboons (Papio). Journal of Human Evolution 146, 102819. (doi:10.1016/j.jhevol.2020.102819)

65. Bergman TJ, Ho L, Beehner JC. 2009 Chest color and social status in male geladas (Theropithecus gelada). International Journal of Primatology 30, 791–806. (doi:10.1007/s10764-009-9374-x)

66. Xia W, Grueter CC, Ren B, Zhang D, Yuan X, Li D. 2021 Determinants of Harem Size in a Polygynous Primate: Reproductive Success and Social Benefits. Animals 11, 2915. (doi:10.3390/ani11102915)

67. Rosenbaum S, Vigilant L, Kuzawa CW, Stoinski TS. 2018 Caring for infants is associated with increased reproductive success for male mountain gorillas. Scientific Reports 8, 15223. (doi:10.1038/s41598-018-33380-4)

68. Goffe AS. 2016 Social relationships of female Guinea baboons (Papio papio) in Senegal. PhD Thesis, Georg-August-Universität Göttingen.

## Supplementary Information References

1. Boese GK. 1975 Social behavior and ecological considerations of West African baboons (Papio papio). In Socioecology and Psychology of Primates (ed Ed Russell H. Tuttle), pp. 205–230. Chicago: De Gruyter Mouton.

2. Dal Pesco F. 2020 Dynamics and fitness benefits of male-male sociality in wild Guinea baboons (Papio papio). PhD Thesis, Georg-August University, Göttingen. See https://ediss.uni-goettingen.de/handle/21.11130/00-1735-0000-0005-134F-E.

3. Dal Pesco F, Trede F, Zinner D, Fischer J. 2021 Kin bias and male pair-bond status shape male-male relationships in a multilevel primate society. Behav Ecol Sociobiol 75, 24. (doi:10.1007/s00265-020-02960-8)

4. Beehner JC, Gesquiere L, Seyfarth RM, Cheney DL, Alberts SC, Altmann J. 2016 Corrigendum to ‘Testosterone related to age and life-history stages in male baboons and geladas’ [Horm. Behav. 56/4 (2009) 472-480]. Hormones and Behavior 80, 149. (doi:https://doi.org/10.1016/j.yhbeh.2015.08.004)

5. Patzelt A, Kopp GH, Ndao I, Kalbitzer U, Zinner D, Fischer J. 2014 Male tolerance and male-male bonds in a multilevel primate society. Proceedings of the National Academy of Sciences of the United States of America 111, 14740–14745. (doi:10.1073/pnas.1405811111)

6. Städele V, Roberts ER, Barrett BJ, Strum SC, Vigilant L, Silk JB. 2019 Male-female relationships in olive baboons (Papio anubis): Parenting or mating effort? Veronika. Journal of Human Evolution journal 127, 81–92.

7. Schielzeth H, Forstmeier W. 2009 Conclusions beyond support: Overconfident estimates in mixed models. Behavioral Ecology 20, 416–420. (doi:10.1093/beheco/arn145)

8. Barr DJ, Levy R, Scheepers C, Tily HJ. 2013 Random effects structure for confirmatory hypothesis testing: Keep it maximal. Journal of Memory and Language 68, 255–278. (doi:10.1016/j.jml.2012.11.001)

9. Matuschek H, Kliegl R, Vasishth S, Baayen H, Bates D. 2017 Balancing type I error and power in linear mixed models. Journal of Memory and Language 94, 305–315.

10. Schielzeth H. 2010 Simple means to improve the interpretability of regression coefficients. Methods in Ecology and Evolution 1, 103–113. (doi:10.1111/j.2041-210X.2010.00012.x)

11. Lüdecke D, Ben-Shachar M, Patil I, Waggoner P, Makowski D. 2021 performance: An R Package for Assessment, Comparison and Testing of Statistical Models. Journal of Open Source Software 6, 3139. (doi:10.21105/joss.03139)

12. Nieuwenhuis R, Grotenhuis M, Pelzer B. 2012 influence.me: Tools for detecting influential data in mixed effects models. The R Journal 4, 38–47.

13. Kitchen DM, Seyfarth RM, Fischer J, Cheney DL. 2003 Loud calls as indicators of dominance in male baboons (Papio cynocephalus ursinus). Behavioral Ecology and Sociobiology 53, 374–384. (doi:10.1007/s00265-003-0588-1)

14. Kalinowski ST, Taper ML, Marshall TC. 2007 Revising how the computer program CERVUS accommodates genotyping error increases success in paternity assignment. Molecular Ecology 16, 1099–1106. (doi:10.1111/j.1365-294X.2007.03089.x)

